# Learning to play with spikes. Characterizing, predicting, and engineering unsupervised plasticity rules for spiking reservoir computing

**DOI:** 10.64898/2026.09.15.751573

**Authors:** Maciej Kania, Basile Confavreux, Tim P. Vogels

**Affiliations:** Institute of Science and Technology Austria; Gatsby Computational Neuroscience Unit University College London

## Abstract

Spiking reservoir computing, and reservoir computing more generally, is a powerful and efficient framework for neuromorphic and biological applications, in which a fixed random reservoir drives a trained readout. Its performance depends critically on the reservoir initialization, so that enriching the reservoir with adaptive, unsupervised plasticity rules offers a natural solution to this limitation. However, it is unclear which rules work, or why, and how to find the best-suited rules for specific tasks without exhaustive and expensive search. Here, we learn how to play the Atari game Pong in a plastic spiking reservoir network. We systematically characterize a large family of local plasticity rules that were meta-learned in prior work. We then show that their performance is predictable and structured: high-scoring rules are characterized by strong differentiation between neurons encoding the ball trajectory and the background, with stable weight dynamics, and consistent readout alignment across time. These mechanistic signatures are not task-specific and transfer to a delayed-match recognition task. Rule performance can be estimated from rule parameters alone. Finally, conditioning simulation-based inference (SBI) on high scores allows us to sample directly from promising regions of the rule space, to discover rules that exceed the performance of those found in the prior distribution and reveal stable, high-performing configurations. Together, these results offer competitive performance against classical reservoir computing while providing a transparent, interpretable account of what makes a plasticity rule useful for neuromorphic hardware and biological computing.

**TL;DR:** Unsupervised local learning rules that stabilize spiking reservoirs also structure their representations – creating interpretable signatures of expertise that enable prediction and engineering of high-performing learning algorithms.

## 1 Introduction

Reservoir computing offers an efficient framework for machine learning: rather than training a complex recurrent network end-to-end, a fixed random reservoir transforms inputs into rich dynamic representations that a simple linear readout learns to decode [1]. This approach significantly reduces the cost and duration of training. However, this efficiency comes at a cost: the reservoir’s representational capacity relies entirely on its fixed initial connectivity and cannot flexibly adapt to a task. This is a fundamental ceiling — readout optimization cannot compensate for a reservoir that fails to represent the task structure. Interestingly, biological networks suggest an alternative — they maintain large recurrent networks that generate complex input representations, while simultaneously allowing plastic changes in their connections [2, 3]. This principle is echoed in the emerging field of biological reservoir computing, most strikingly in brain organoid computing, where lab-grown neural networks presumably preserve learning mechanisms that organize network dynamics and allow for solving complex tasks [4–6]. Yet which specific learning mechanisms enable these computations remains an open question.

Spiking neural networks are a natural substrate for reservoir computing, offering both biological plausibility and energy efficiency compatible with neuromorphic hardware. Yet how unsupervised plasticity shapes their dynamics remains far from being understood. Intriguingly, the brain putatively implements a spectrum of learning mechanisms, from unsupervised, local plasticity rules [7] to global reinforcement signals [8, 9], representing an enormous space of possible solutions. From an engineering perspective, this space is both a challenge and an opportunity: it is too large to search exhaustively, but it may contain plasticity rules that are excellent for neuromorphic or reservoir computing systems. Crucially, while such solutions differ across systems and functions, they share one important characteristic: degeneracy [10–12] — meaning that many different rule configurations could possibly produce equivalent, high-performing outcomes. This principle has recently emerged prominently in activity-dependent synaptic plasticity, where multiple research groups discovered families of learning rules: rules that stabilize network dynamics and produce memory as a byproduct [13, 14], rules that maintain circuit function under perturbation [15], and rules that organize multi-cell-type activity across cortical layers to perform visual tasks [16]. This suggests that highly-functional plasticity rules are not rare flukes but occupy structured, discoverable regions of parameter space.

Here, we aim to investigate whether biologically constrained plasticity rules can serve as unsupervised local learning algorithms for reservoir computing. Specifically, we aim to understand what makes a plasticity rule effective for reservoir computing, how to predict its performance from mechanistic signatures, and how to engineer new, better algorithms. As a testbed, we use the Atari game Pong [17], a task with well-defined temporal structure and variable ball dynamics, and further test whether the mechanistic signatures we identify generalize beyond it to the delayed-match task. We systematically screen the full population of rules discovered in prior work [13] as potential learning algorithms for reservoir computing. Prior work in rate-coded reservoirs has linked plasticity-induced performance gains to changes in reservoir activity structure [18]. Whether the activity-structure signatures found in rate-coded reservoirs extend to biologically realistic spiking reservoirs, and whether they can guide active rule engineering, is the focus of this study.

We focus on assessing the contribution of the reservoir dynamics and readout features to the performance of the model and linking the structure of the rule to its functionality. We find that high-scoring models form stable representations of the task, driving a contrast between neurons encoding the current and previous ball positions and the background, while low-scoring models create weaker contrast and drift more. These signatures let us predict performance from rule parameters alone (computation-free), and, using simulation-based inference [19–23], sample high-score regions of rule space to recover rules that outperform exhaustive screening.

## 2 Enriching reservoir computing with local unsupervised plasticity rules in the Pong framework

We start by adopting a complex reservoir computing task, the Atari game Pong [17, 24] (Fig. 1). Prior work [13] used the Pong task to show 3 examples of plasticity rules that, largely through offline training, learned to predict the direction of the ball. Here, we expand on these experiments and screen the entire family [25] of stable plastic networks in a more rigorous, online training paradigm.

**Figure 1:**
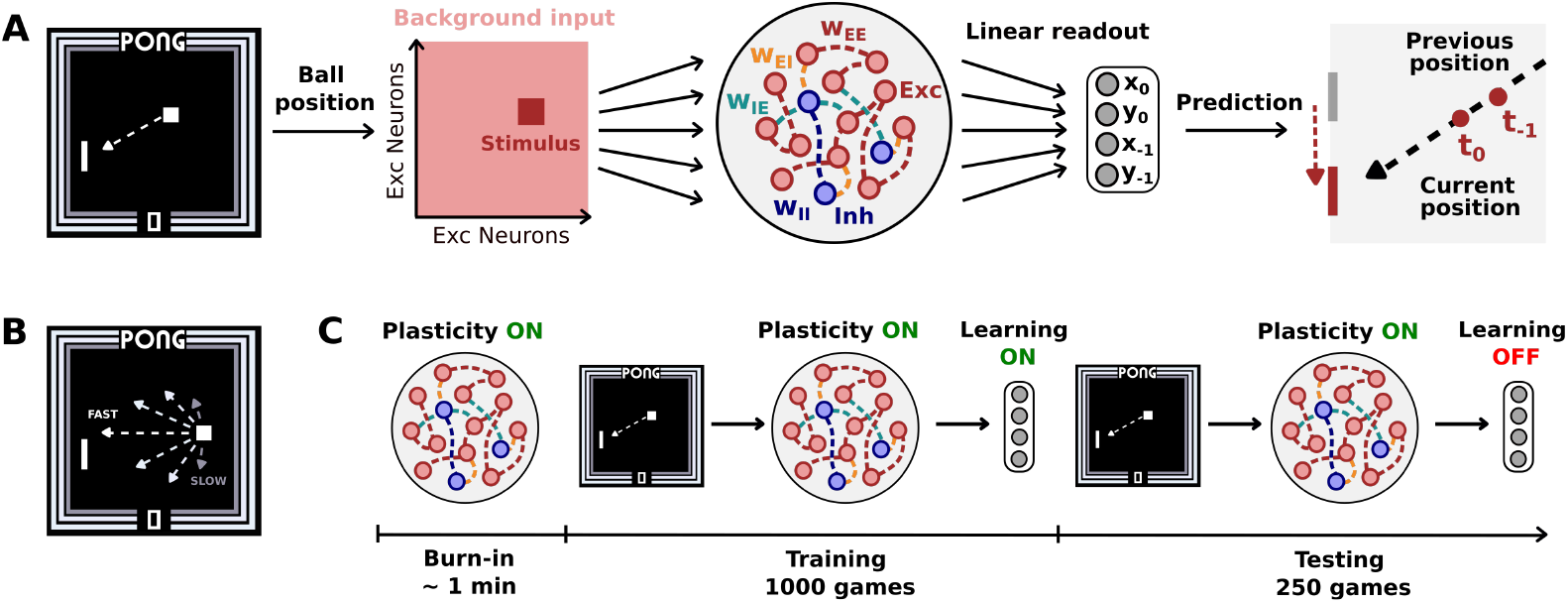
Reservoir computing framework for biologically-plausible unsupervised learning in Atari Pong. **A**. Ball position is encoded as spatial input to a recurrent spiking network of excitatory and inhibitory neurons; a linear readout predicts the ball’s direction based on its current and previous positions. **B**. The task includes variable ball speeds and angles. **C**. Training progresses in three steps: a 1-minute burn-in, 1000 training games, and 250 testing games (Technical appendix).

We keep the same Pong implementation: ball position drives a spatial input to a spiking reservoir, and a linear readout decodes ball direction to move the paddle (Fig. 1A). In each game, the ball direction is randomly initialized, and its velocity may change after a contact with the walls, making prediction less trivial (Fig. 1B). Unlike the original work, we begin an online learning paradigm immediately following the network’s burn-in period (Fig. 1C). In the previous work [13], the network was trained by replaying simulated games all rules saw the same number of ball trajectories during the training. By introducing online learning here, we explicitly take advantage of the Matthew’s effect [26]: If a plastic model is good at playing, games last longer, and the total number of ball trajectories encountered during training is higher than for rules that frequently fail to hit the ball. This design couples rule quality with training exposure. We control for this coupling directly with a fixed-trajectory protocol that equalizes the number of ball trajectories seen by every rule (Fig. S1); we otherwise adopt online learning to mirror the conditions under which biological and neuromorphic systems operate, where better-performing networks naturally accumulate more structured input.

### 2.1 Evaluation of the family of activity-dependent plasticity rules in Pong

We begin by implementing a recurrent spiking neural network comprising 4096 excitatory (E) and 1024 inhibitory (I) leaky-integrate-and-fire (LIF) neurons as a reservoir network with random, sparse (10%) connectivity. In contrast to classical reservoir computing, the synaptic weights follow activity-dependent updates according to plasticity rules formalized as:

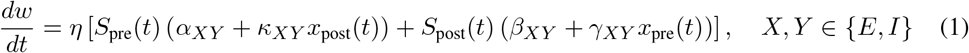

where *S*_pre_(*t*) and *S*_post_(*t*) are spike trains, and *x*_pre_(*t*) and *x*_post_(*t*) are their low-pass filtered versions (Technical appendix). A single plasticity rule is specified by six parameters for each of the four synapse types *XY* ∈ {*EE, EI, IE, II*}: the four amplitudes(*α, β, γ, κ*) of Eq. (1), where *α, β* are trace-independent baselines and *κ, γ* weight the coincidence of spiking with the partner’s filtered activity, and two timescales(*τ*_pre_, *τ*_post_), giving a 24-dimensional parameter vector (Technical appendix, Eqs. 5–6). We evaluate a fixed library of 2201 such rules inherited from prior work [13, 21], where they were meta-learned to produce stable, cortical-like activity. Each rule is evaluated across three seeds (Fig. 2A).

**Figure 2:**
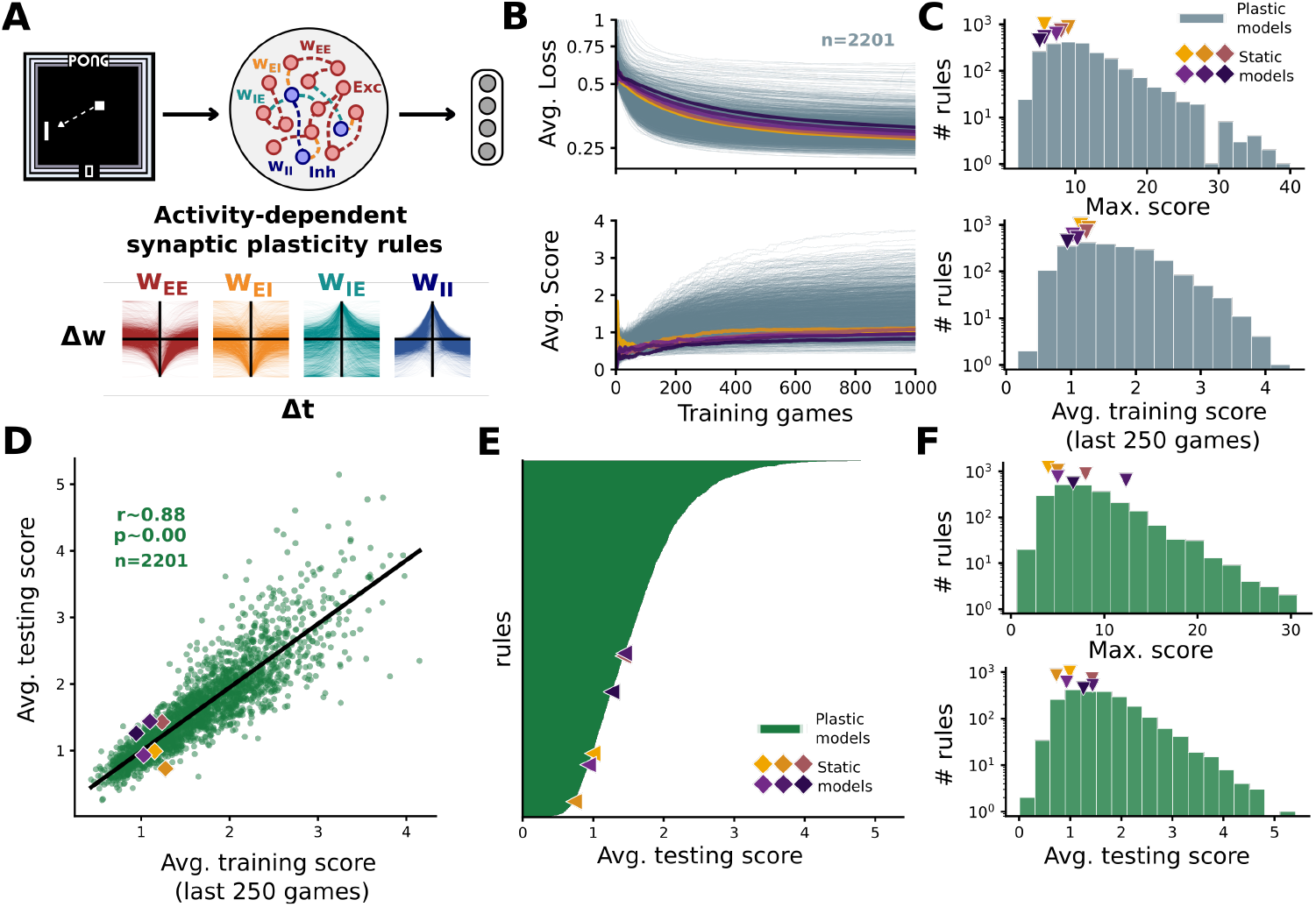
The family of learning rules exhibits a wide range of expertise in Pong, outperforming static reservoir networks. **A**. Combination of four activity-dependent synaptic plasticity rules (*w*_*EE*_, *w*_*EI*_, *w*_*IE*_, *w*_*II*_) governing reservoir dynamics. **B**. Average loss and score across 1000 training games for plastic models in grey, with static models in yellow and purple. **C**. Distributions of maximum scores over 1000 games and training scores in the last 250 games averaged across three seeds. **D**. Correlation between average training score (last 250 games) and average testing score across all plastic models. **E**. Plastic models ordered by the average score during testing. **F**. Distributions of maximum scores and testing scores averaged across three seeds.

The performance of each rule is evaluated in a stereotyped protocol (Fig. 1C) that features a minute long equilibration (burn-in) period, followed by 1000 training games. During the training phase, all plastic models (in grey) and models without plasticity with varying levels of connectivity heterogeneity (referred to as static, from yellow to purple, Technical appendix) decreased their loss and increased their average score (Fig. 2B). Notably, we observe a wide range of expertise, from rules that produced performance far below that of static networks, to expert rules that reliably gained scores during the games (Fig. 2C). Performance achieved during the testing phase was highly correlated with the score during the last 250 games of the training phase (r=0.88, p<0.001, n=2201), confirming a wide range of performance across the family of plasticity rules (Fig. 2D). The distribution of scores was highly skewed, with the majority of rules having low-to-moderate performance, below 2 points on average per game (Fig. 2E-F). Interestingly, when ordered by performance, we observe a sharp increase in performance for the 5-10% of the rules (about 100-200 rules score above 3 points on average per game). These may serve as suitable learning algorithms for reservoir computing. Under the fixed-trajectory protocol, the rule-quality ordering was preserved: per-rule scores under equalized exposure correlated strongly with the original closed-loop scores across all screened rules (Spearman *ρ* = 0.84; Fig. S1). The Matthew-effect coupling therefore amplifies, but does not create, the differences between rules. After characterizing the plastic models’ performance and identifying expert-like rules, we seek to understand what underlies this range of function across the models.

## 3 Disentangling the contributions of rules, reservoir, and readout features to Pong performance

Predicting how well a given learning rule performs in a task solely on its parameters would allow precise identification of a small group of expert-like candidate models. However, previous studies found no such relationship. While predicting the activity of excitatory and inhibitory populations came with ease (r^2^≈0.7), predicting the longevity of the learning rule or its memory function solely on its parametrization remains unreliable (r^2^ ≈ 0.2 and 0.1, respectively) [13, 14]. Instead, we exploit our reservoir computing setup to identify what separates highfrom low-scoring models across readout weights, reservoir activity, and weight dynamics, and then proceed to predict the performance.

### 3.1 Decomposing readout, activity, and weight features of high- and low-scoring models

First, we select the highest-, mid-, and lowest-scoring models, each comprising 5% (111 rules) of all screened rules (Fig. 3A–B). As expected, better-performing rules show lower average loss at the end of training (Fig. 3C). The average testing score was strongly correlated with average loss in log-log space across all screened rules (r=-0.77, p<0.001, n=2201; Fig. 3D). We next examine the contribution of each component (readout weights, reservoir activity, and synaptic weights) to model performance.

**Figure 3:**
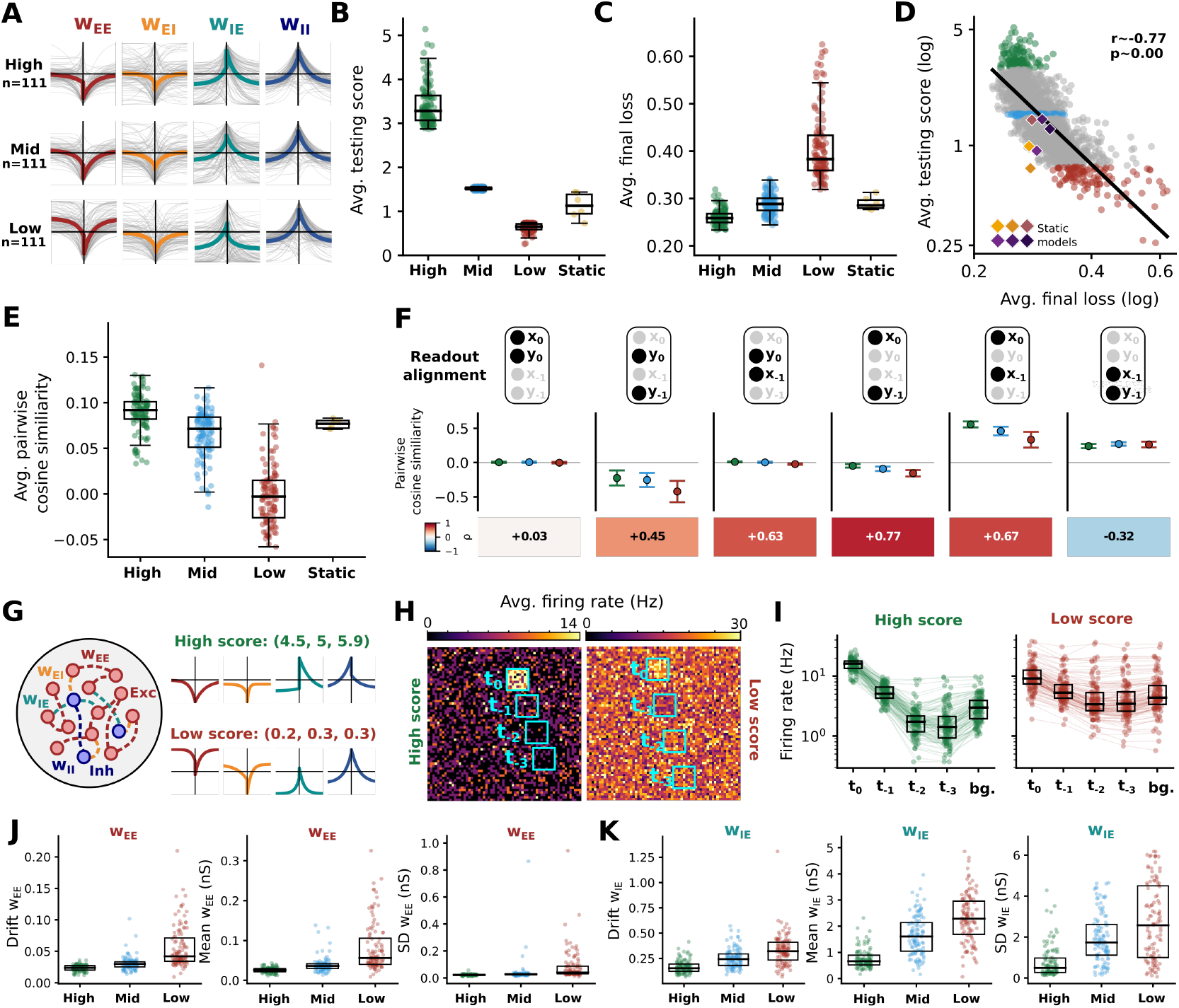
Models that better predict ball direction also show better-aligned readout units and stable and more structured spatial representations of the stimulus. **A**. High-, mid-, and lowperforming 5% of all screened plastic models. Individual rules in gray, average shape of the rule in colors. **B**. Average testing score and **C**. average loss after training across high, mid, low, and static models. **D**. Log-log correlation between average loss after training and average testing score across all plastic models. **E**. Average pairwise cosine similarity of readout weights across plastic models. **F**. Pairwise cosine similarity between readout units encoding current and previous ball positions, with correlation values between them and average testing score. **G**. Example of high and low performing models. **H**. Average firing rate of excitatory neurons in the reservoir high-scoring (left) and low-scoring (right) networks arranged to match the space of the game. Blue squares represent current (*t*_0_) and previous positions (*t*_*−* 1_, *t*_*−* 2_, *t*_*−* 3_) of the ball. **I**. Firing rates of excitatory neurons at the current and previous ball positions (*t*) and the rest of the population (*bg*.) **J–K**. Drift, mean, and standard deviation (SD) of synaptic weights *w*_*EE*_ and *w*_*IE*_ across high, mid, and low performing models.

#### Readout

We start by analyzing the readout, as it directly affects how well a plastic model predicts the ball direction. Our readout has four units, each predicting a different coordinate for present (*x*_0_, *y*_0_) or past (*x*_*−* 1_, *y* _*−* 1_) positions of the ball (Technical appendix). We find that high-scoring models on average across the pairs of units tend to align their weight vectors more than worse-performing models (Fig. 3E). We assess this in detail by separately looking at each pair. The alignment of units that both decoded coordinates in either past or present had no or weak negative correlation with the performance of the model (Fig. 3F, first and last panel). However, units that decode coordinates across two different time points tend to be more aligned or closer to orthogonal in high-scoring models, in contrast to the low-scoring ones (Fig. 3F, four middle panels). Finally, alignment across all screened rules for the pairs decoding across time points positively correlated with the average testing score (Fig. 3F, bottom boxes, n=2201).

#### Reservoir activity

The linear readout decodes the ball’s direction based on the activity of the excitatory population (4096 neurons). We look deeper into the dynamics of the high- and low-scoring networks during the Pong task (Fig. 3G-H). We observed that the high-scoring model had a clear contrast between the neurons receiving spatial input (*t*_0_) and background neurons (bg), while the stimulated neurons in the low-scoring model blended with the background activity. Interestingly, by looking at the past positions of the ball (*t*_*−*1_, *t* _*−*2_, *t* _*−*3_), we observe a decrease in activity below the background rate, which reveals the ball trajectory within a single time frame. We quantify and confirm this for 111 high- and 111 low-scoring models (Fig. 3I).

#### Reservoir weights

Finally, the dynamics of the reservoir are a product of network connectivity. Here, we focus on the excitatory-to-excitatory (*w*_*EE*_; Fig. 3J) and inhibitory-to-excitatory (*w*_*IE*_; Fig. 3K) synaptic strengths after training. We computed the difference between the weights after training and after testing. High-scoring models had smaller and more stable weights, while the weights of the worst models were larger and drifted more. This suggests that the best models establish stable representations of the ball trajectory during training, while the low-scoring ones continue changing.

### 3.2 Pong score prediction based on the signatures of learning

In Section 3.1, we find good predictors of the learning rule performance in Pong. However, these features are available only after full training, which for the expert models is computationally expensive. Here, we prepare three levels of linear regression models - computation-free (Tier 1), computationally cheap (Tier 2), and full computation (Tier 3) (Table 1).

**Table 1:** Ridge regression models predicting average testing score from different features of plastic models.

|  | Regression model |  | *k* | CV *r*<sup>2</sup> | CV RMSE |
|---|---|---|---|---|---|
| Single | **Tier 1** | **Plasticity rule parameters** | 24 | **0.352** | 0.549 |
|  | Tier 2a | Novelty detection | 10 | 0.125 | 0.638 |
|  | Tier 2b | Slope of the loss during training | 1 | 0.338 | 0.555 |
|  | **Tier 2** | **Novelty detection + Loss slope** | 11 | **0.353** | 0.548 |
|  | Tier 3a | Readout alignment | 12 | 0.648 | 0.404 |
|  | Tier 3b | Activity contrast | 9 | 0.619 | 0.421 |
|  | Tier 3c | Reservoir weights | 20 | 0.380 | 0.537 |
|  | **Tier 3** | **Readout + Activity + Weights** | 41 | **0.722** | 0.359 |
| Combined | Tier 1+2 |  | 35 | 0.529 | 0.468 |
|  | Tier 1+3 |  | 65 | 0.720 | 0.360 |
|  | Tier 2+3 |  | 52 | 0.726 | 0.357 |
|  | **Tier 1+2+3** |  | 76 | **0.723** | 0.359 |

In **Tier 1**, we predict model performance solely from the structure of the learning rules (24 parameters). Even with just the shape of the plasticity rule, we obtain a modest predictive power (r^2^ 0.35; details in Technical appendix). Although not fully satisfactory, this represents a notable improvement over previous works [13, 14]. Allowing a nonlinear predictor (TabPFN [27]) raises parameter-only prediction substantially (*R*^2^ ≈ 0.56 vs. linear *R*^2^ ≈ 0.35; Table S1), indicating the 24-dimensional rule parametrization carries more performance-predictive signal than a linear model captures. In **Tier 2**, we add features that can be computed cheaply: novelty detection requiring only 1–2 min of simulation (Technical appendix) and/or the initial slope of the loss during training (accessible within 200 games). Tier 2 alone yields similar predictive power (r^2^ ≈ 0.35); however, combining Tiers 1 and 2 results in moderate-to-high prediction (r^2^ ≈ 0.53). Finally, **Tier 3** (full training) combines readout alignment, activity contrast, and reservoir weight features, yielding strong predictive power (r^2^ ≈ 0.72). Notably, combining all three tiers did not further improve prediction (r^2^ ≈ 0.72).

## 4 Generalization of mechanistic signatures to a delayed-match task

To test whether the signatures of expertise we identified are specific to Pong or reflect a general property of the plasticity rules, we evaluated the same plastic models on a structurally different task: a delayed-match (DM) recognition task (Fig. 4A). On each trial, two stimulus patterns drawn from a pool of 13 (non-overlapping) assemblies are presented sequentially, separated by a 5 s delay, and a linear readout reports whether the two patterns match.

**Figure 4:**
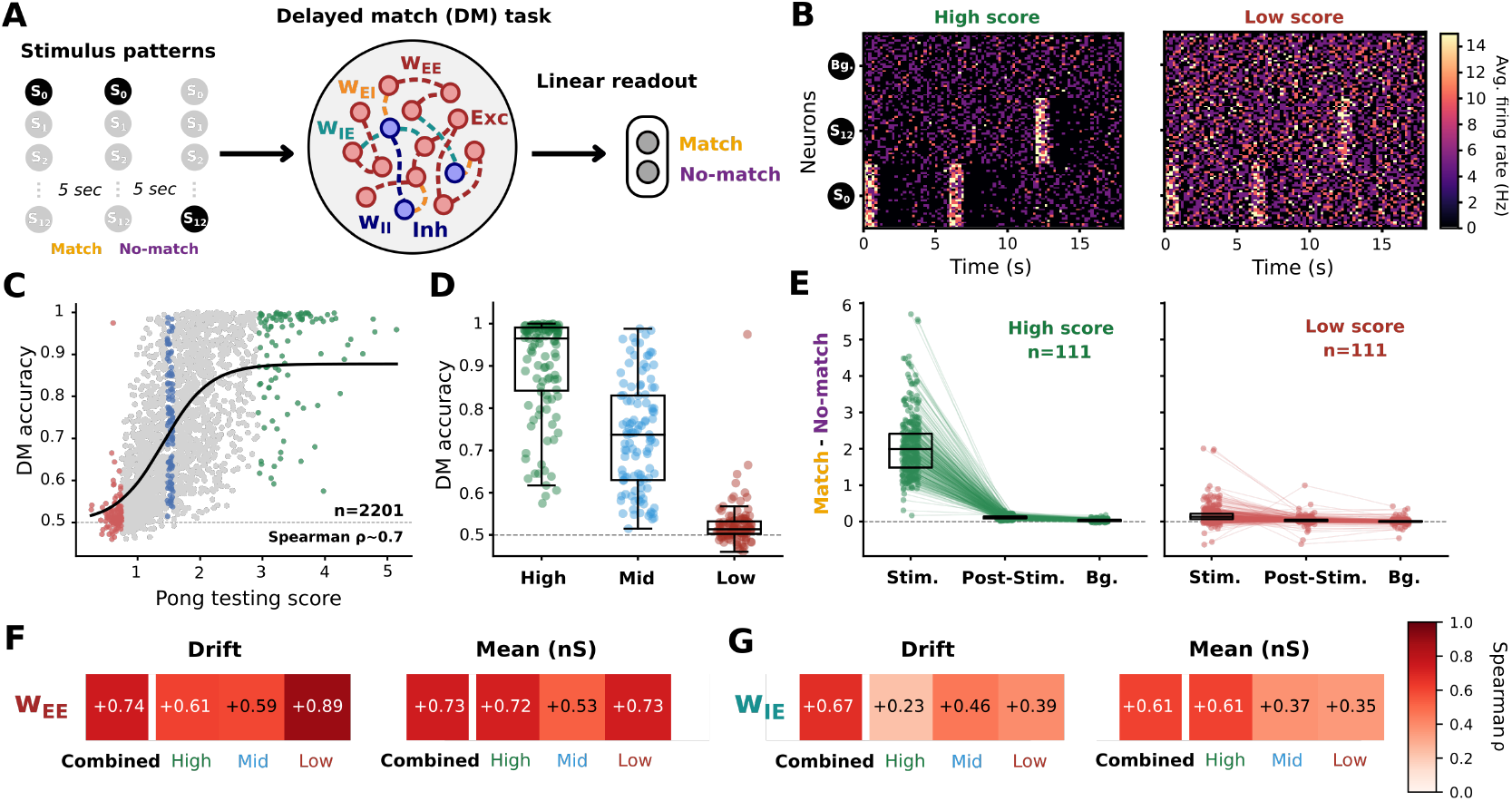
Generalization to a delayed-match (DM) task. **A**. DM task schematic. On each trial, two stimulus patterns are presented sequentially to the network, separated by a 5 s delay. A linear readout then reports whether the two patterns match or not. **B**. An example high- and low-scoring rule, with neurons grouped by assembly (*S*_0_, *S*_12_, background) and aligned to stimulus onset. **C**. DM accuracy vs. Pong testing score across all screened rules (n=2201), with a logistic fit and points coloured by Pong score rank (high/green, mid/blue, low/red) (Spearman *ρ* ~ 0.7); chance is 0.5 (dashed). **D**. DM accuracy for the high-, mid-, and low-scoring Pong ranks. **E**. Match vs. no-match response for high- (n=111) and low-scoring (n=111) rules during the stimulus (Stim., 1 s), post-stimulus (Post-Stim., 5 s), and background (Bg., 6 s) windows. **F–G**. Cross-task Spearman *ρ* between the Pong and DM weight signatures for *w*_*EE*_ (**F**) and *w*_*IE*_ (**G**), for drift and mean weight (nS).

Overall, DM task performance tracked the Pong expertise (Fig. 4C). Across all screened rules (n=2201), DM accuracy increased with Pong testing score following a saturating relationship (Spearman *ρ* ~0.7; chance = 0.5) and the high-, mid-, and low-scoring Pong models could be cleanly separated (Fig. 4D). The sigmoidal relationship between performance in those tasks suggests that being a good Pong model is thus sufficient but not necessary for DM expertise: above a threshold level of Pong expertise, rules reliably support the DM task. We then further investigated the mechanistic features during the DM task. High-scoring rules produced sharp, assembly-specific responses that were absent in low-scoring rules, whose activity blended into the background (Fig. 4B), which mirrored the activity-contrast signature we found in Pong. Quantifying the difference between assembly responses during the match and non-match conditions, we found that high-scoring models produced on average stronger responses during the match conditions, while low-scoring models had almost no difference between those two cases (Fig. 4E). Finally, the weight signatures transferred across tasks. Per-rule *w*_*EE*_ and *w*_*IE*_ drift and mean weight were strongly correlated between Pong and DM (Spearman *ρ* = 0.74 and 0.73 for *w*_*EE*_; *ρ* = 0.67 and 0.61 for *w*_*IE*_; Fig. 4F-G). Together, these results show that activity contrast and weight dynamics are not Pong-specific but general markers of unsupervised plasticity models.

## 5 Benchmarking against other reservoir computing models

So far, we have compared the learning rules with one another and with their static equivalents. To further evaluate whether these learning rules can serve as an alternative to classical reservoir computing and neuromorphic approaches, we benchmarked the best plastic rules, corresponding to the top 5% of screened rules (n=111), against classical reservoir computing setups and algorithms for training spiking networks (Fig. 5). For each baseline we report the plastic budget *k*^*∗*^ at which the expected best-of-*k* rule matches its best score, keeping the comparison budget-fair (Table S2). For details on each configuration and its performance, see the Technical appendix.

**Figure 5:**
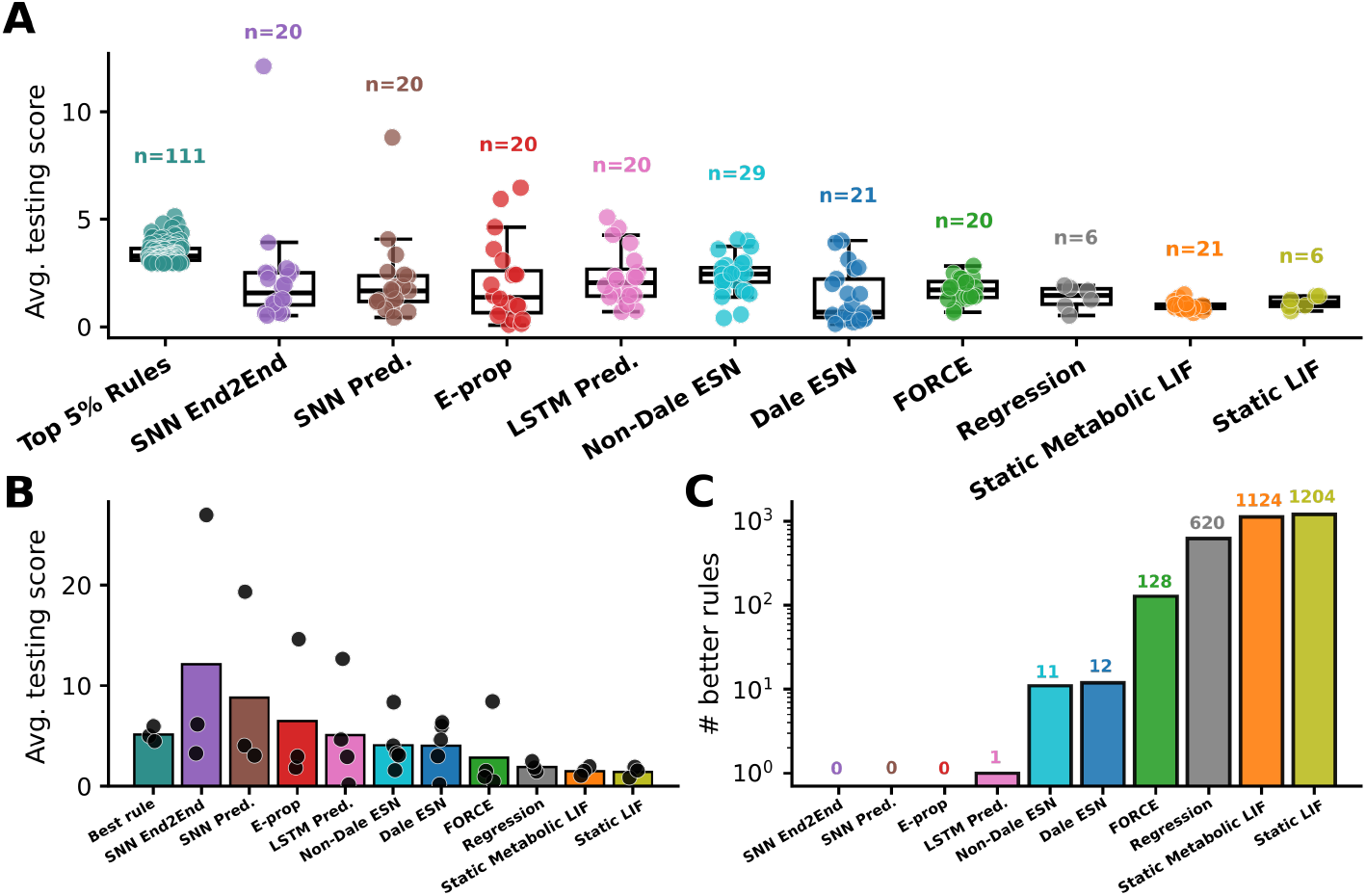
Benchmark of the best plastic learning rules against classical learning models. **A**. Distribution of average testing scores across the top 5% of screened plastic rules (n=111) and representative configurations of baseline models: surrogate-gradient SNNs trained end-to-end (SNN End2End) and predictively (SNN Pred.), e-prop, LSTM trained predictively (LSTM Pred.), echo state networks without (Non-Dale ESN) and with Dale’s law (Dale ESN), FORCE learning, linear regression without a reservoir, and static spiking networks with and without a metabolic homeostatic process. **B**. Average testing score of the single best configuration from each model class. Black dots indicate different initialization seeds for each model. **C**. Number of plastic learning rules that outperform the best configuration of each baseline model class.

We tested multiple parameter configurations across 10 model classes, ranging from non-spiking static reservoirs to fully trained spiking neural networks (SNNs) (Fig. 5A). All models were implemented in the Pong framework with reservoir and readout components, matching our original setup (Fig. 1). We then compared the best-performing configuration within each model class based on its average testing score (Fig. 5B). The best plastic model substantially outperformed its static counterparts, the static SNNs with metabolic homeostasis [28], and the regression-only baselines with varying noise levels. The plastic model also outperformed classical echo state networks (ESNs) [29–31], static reservoirs trained with the FORCE algorithm [32], and fully trained predictive long short-term memory (LSTM) models [33, 34]. However, for these models, the advantage was more modest: only a small subset of plastic rules, ranging from 1 to 128, achieved better performance than the best configuration in each class (Fig. 5C).

In contrast, the best plastic model was competitive with, but did not outperform, SNNs fully trained using e-prop [35] or surrogate gradient descent [36, 37]. Notably, these fully trained SNN models showed a wide spread in performance across initialization seeds, whereas the best plastic model remained comparatively consistent across seeds.

## 6 Engineering expert algorithms via simulation-based inference

Finally, we want to demonstrate that the family of learning rules can serve as a substrate for engineering new rules that achieve expertise in the Pong task (Fig. 6). The family of rules used here was previously discovered using a simulation-based inference (SBI) approach — a Bayesian inference framework for stochastic models whose likelihood cannot be tracked analytically but can be estimated through simulation [19, 21, 23].

**Figure 6:**
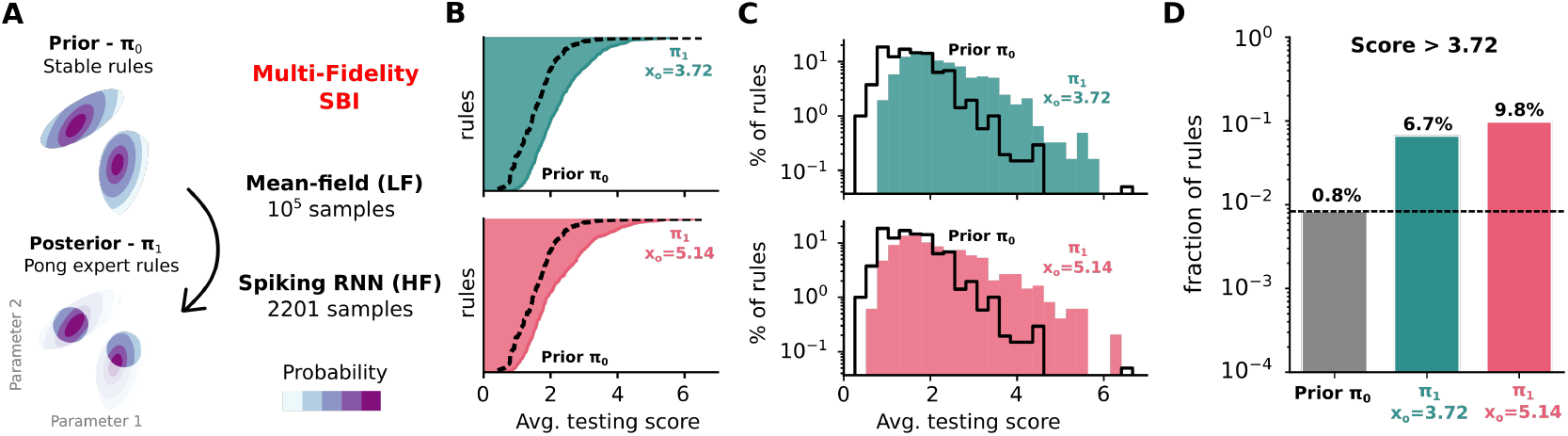
Inferring expert learning rules via multi-fidelity simulation-based inference (SBI) [22]. **A**. Schematic of the MF-SBI pipeline. A mean-field model (low-fidelity) pre-trains the density estimator, which is then refined using the full spiking RNN (high-fidelity). **B**. Average testing scores of rules sampled from the posterior *π*_1_ (green n=615; red n=489), ordered by performance, compared to reweighted subsampled prior rules (*π*_0_, black dashed). **C**. Histograms of average testing scores for rules sampled from *π*_0_ and *π*_1_. **D**. Fraction of rules exceeding a score of 3.72 for the prior *π*_0_ and both posteriors.

Here, we go a step further and ask whether we can discover rules that are experts in our Pong task (Fig. 6A). To this end, we use all screened plastic models as samples drawn from the prior (*π*_0_), which contains rules that result in stable, cortical-like activity. However, given the relatively small sample size (n=2201) and the high dimensionality of the learning rules (24 parameters), we turn to a novel multi-fidelity SBI approach (MF-SBI) [22]. We generate thousands of computation-free mean-field approximations (low-fidelity models, LF) to assess learning rule stability. To avoid evaluating these rules in Pong, we assign each a score predicted by a gradient-boosting surrogate trained on the 2201 previously tested high-fidelity rules (Technical appendix). This allows us to pre-train the density estimator network before performing full training on the smaller set of high-fidelity (HF) models.

We sample and simulate new learning rules from the posterior (*π*_1_) using two observations: the maximum testing score across prior samples (*x*_*o*_=5.14, n=900) and the average score of the top 1% of previously screened rules (*x*_*o*_=3.72, n=934). While the previous work [21] conditioned the prior for stability, it does not ensure that newly sampled rules will maintain stable, cortical-like activity during the Pong task. We therefore apply a stability proxy and filter out rules whose firing rates fall outside 1–50 Hz or whose weights blow-out by the end of training (see Technical appendix). We find that both filtered samples (*x*_*o*_=3.72; *x*_*o*_=5.14) contain more high-scoring models than the prior distribution (Fig. 6B–D). To confirm this enrichment does not merely reflect the SBI-conditioning seeds, panels B–D re-score all rules on three held-out seeds; per-rule scores are preserved and both posteriors stay enriched relative to the reweighted prior (Fig. S2).

## 7 Discussion

In this study we characterize, understand, and engineer plasticity rules that improve spiking reservoir computing, using Atari Pong as a testbed. We found that a large solution space of learning rules comes with a wide range of expertise, including a handful of high-scoring plastic models. We showed that high-scoring rules can be clearly separated from the low-scoring ones based on the learning rules parametrization and their general dynamical properties. Most importantly, we show that features of readout and reservoir learning strongly separate those models - expert rules drive structured and stable representations of the input, while the low-scoring models had little contrast between the activity of the background neurons and those encoding ball trajectory, and their weights showed a high level of drift.

What kind of learning drives expert performance? The plasticity rules we use are designed to encode long-term changes through persistent synaptic weight updates. However, when a freshly burned-in reservoir is paired with a readout trained on a fully plastic network, it still achieves moderate-to-high performance (see the Supplementary figure S3), suggesting that short-timescale dynamics, resembling activity-silent working memory [38], may also contribute, transiently retaining the ball trajectory in network activity. At the same time, freezing the reservoir after training still yields high performance (see the Supplementary figure S4), confirming that long-term synaptic reorganization is genuinely taking place. These results suggest that unsupervised plasticity rules may drive expert performance through a mixture of working-memory-like dynamics and long-term changes in connectivity.

Our work builds on a growing body of research showing that unsupervised local plasticity rules can improve reservoir computing beyond static random connectivity. In rate-coded echo state networks, prior work established that plasticity-induced performance gains are linked to reduced pairwise correlations and increased input separability [18], that heterogeneous rules discovered via evolutionary search can further boost performance [39], and that growing task-specific connectivity from scratch via unsupervised Hebbian rules offers a complementary structural route [40]. A related line of work implements structural plasticity through dynamic sparse training guided by topological link-prediction rules (epitopological learning) [41, 42]. Our findings extend this picture to a biologically realistic spiking substrate with E/I populations, decomposing the mechanistic signatures into activity contrast, readout alignment, and weight stability — and showing that these signatures can be used not just descriptively but predictively, to engineer better rules via SBI. On the spiking side, early work showed that STDP combined with intrinsic plasticity produces fading memory and structured dynamics in recurrent spiking networks [2, 3], consistent with our finding that high-performing rules drive stable, structured representations. Relative to this literature, our work proposes a complete loop from mechanistic understanding to active engineering of learning rules for reservoir computing.

## Possible applications

The learning rules operate locally and unsupervised, making them natural candidates for *on-chip learning in neuromorphic hardware* with explicit E/I architectures [43–46], with the possibility of pretraining weights offline and transferring them to a fixed spiking chip. On the other hand, our work goes in parallel with advances in organoid computing, which realizes reservoir computing in living neural systems [4–6]. We see plastic recurrent spiking networks as natural *digital twins* [12] for those: constrained to match organoid properties (neuron count, E/I ratio, connection sparsity, and activity statistics) they could enable long-duration simulations, systematic parameter sweeps, and protocol testing before translation to *in vitro* neuronal cultures that are fragile and experimentally demanding.

## Limitations

We evaluate the plastic models under two tasks and find that the same mechanistic signatures govern both. However, the two tasks share the same reservoir, assembly structure, and activity-contrast mechanism, so they probe a limited task family rather than establishing taskgeneral signatures. Testing on a structurally distinct benchmark, such as the Heidelberg Spoken Digits dataset [47], a standard temporal classification task for spiking networks, would establish whether activity contrast, readout alignment, and weight stability are universal signatures of effective unsupervised plasticity or task-specific ones. We maintained the reservoir size and readout architecture of previous work [13] to enable direct comparison, but note that a lower-dimensional readout would more cleanly isolate the reservoir’s contribution to performance.

Finally, our plasticity rules operate on short timescales of 100–200 ms and follow an unsupervised, activity-dependent character. The neuroscience of learning ranges across many additional mechanisms, including behavioural timescale synaptic plasticity which operates on the scale of seconds and carries a more instructive, supervised character [48, 49], as well as reward-modulated plasticity rules that give rise to reinforcement learning [8], as well as structural (epitopological) plasticity, which reshapes connectivity rather than only synaptic weights and could be co-adapted with our fixed-topology reservoir via dynamic sparse training [41, 42]. Beyond reshaping an existing topology, the initial connectivity itself could be structured rather than random, for instance using dendritic or receptive-field-based initialization [50] instead of the random sparse connectivity we adopt. Exploring whether such mechanisms further improve spiking reservoir performance remains an exciting direction for future work.

To sum up, our work shows that the performance of unsupervised plasticity rules in spiking reservoir networks is far from random — it is structured, predictable, and can be actively exploited to engineer better algorithms.

## Acknowledgments and Disclosure of Funding

We would like to thank Chaitanya Chintaluri, Douglas Feitosa Tomé, Zoe Harrington and other Vogels group members, as well as Henning Sprekeler for insightful discussions and feedback. We also thank Nastya Krouglova and Pedro Gonçalves for the training and support. M.K. was funded within the DOC program of the OeAW. This work was funded by the European Research Council (ERC consolidator grant SYNAPSEEK). This research was supported by the Scientific Service Units (SSU) of ISTA through resources provided by Scientific Computing (SciComp). The authors declare no competing interests.

## A Technical appendices and supplementary material

The code and data to reproduce the results in this paper can be found at: https://github.com/VogelsLab/spiking-pong. Spiking network simulations were performed using the Auryn simulator [51]. The screening of plasticity rules (Fig. 2), the benchmark (Fig. 5), and simulation of new, expert rules (Fig. 6) were performed on 1152 CPUs on the HPC cluster. Each rule evaluation used 2 CPUs and less than 1 GB of RAM, and took approximately 1–2 days depending on performance, as high-scoring rules accumulate more games and therefore run longer. Random seeds used for each plasticity rule are reported in the accompanying data file. Pong implementation followed the code from Strong et al. [24] and training of the models was performed with PyTorch [52]. Multi-fidelity SBI was performed following fSBI and MF-SBI repositories [21, 22]. Analysis of all simulations was performed using numpy [53], matplotlib [54], SciPy [55] and scikit-learn [56].

### A.1 Recurrent spiking network model

#### A.1.1 Network model

We considered a recurrent spiking network with *N*_E_ = 4096 excitatory neurons and *N*_I_ = 1024 inhibitory neurons (leaky-integrate and fire point neurons with variable threshold, AMPA and NMDA currents, and conductance-based synapses). This network was based on previous work [57, 13]. The 4:1 excitatory-to-inhibitory ratio matches cortical anatomy (~ 80% of neocortical neurons are excitatory) rather than being a purely engineering choice; the absolute counts are powers of two chosen for computational convenience. The ratio, together with the synaptic weights, sets the dynamical regime placing the network in the asynchronous-irregular regime we rely on for rich reservoir dynamics. The plasticity rules underlying our library were shown to remain within their stability bounds at 3:1 and 5:1 ratios [21], so the framework does not depend on the precise value.

The membrane potential dynamics of neuron *j* (excitatory or inhibitory) followed:

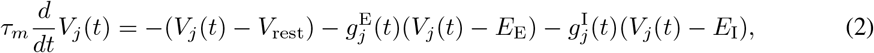

with *τ*_*m*_ = 20 ms, *V*_rest_ = −70 mV, *E*_E_ = 0 mV and *E*_I_ = −80 mV.

A postsynaptic spike occurred whenever the membrane potential *V*_*j*_(*t*) crossed a threshold 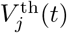, with an instantaneous reset to *V*_reset_ = −70 mV. This threshold 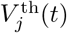was incremented by 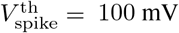 every time neuron *j* spiked and otherwise decayed following:

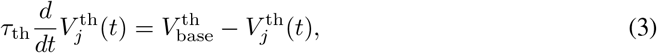

with 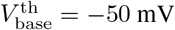. The excitatory and inhibitory conductances, *g*^E^ and *g*^I^ evolved such that

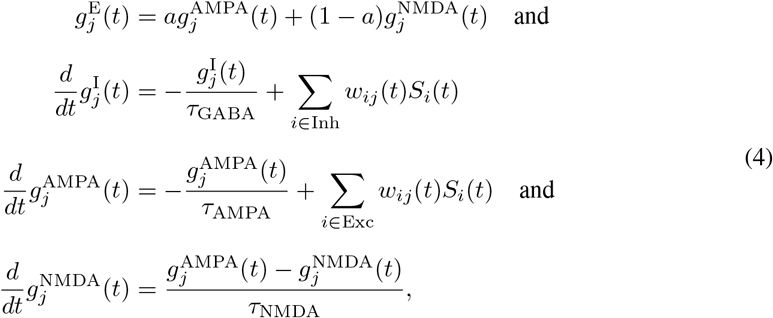

with *w*_*ij*_(*t*) the connection strength between neurons *i* and *j* (unitless), *a* = 0.3 (unitless), *τ*_GABA_ = 10 ms, *τ*_AMPA_ = 5 ms, *τ*_NMDA_ = 100 ms, 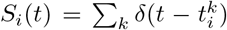 the spike train of presynaptic neuron *i*, where 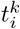denotes the spike times of neuron *k*, and *δ* the Dirac delta.

The network was initialized with random sparse connectivity (10%), with 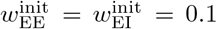 and 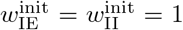. Sparse connectivity at this level is biologically motivated and follows the framework we build on [13]. Because connection density is compensated by weight scaling, the operating regime is preserved across a range of densities; the rules were shown to remain within their stability bounds when connectivity was halved to 5% [21].

The excitatory and inhibitory neurons in the network received *N*_inp*→*E_ = 10000 inputs from Poisson neurons firing at 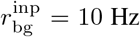. The connectivity from input neurons to excitatory and inhibitory neurons was receptive-field-like: for each recurrent neuron, we selected a random input neuron as the center of the circular receptive field of radius 8. The connections from neurons of this circular patch of input neurons to the considered recurrent neuron was *w*_inp_ = 0.075, and 0 to all other input neurons.

#### A.1.2 Plasticity parameterization

This parameterization of plasticity rules included variations of spike-timing-dependent plasticity, and was taken from previous work [21, 13]. The weight from neuron *i* to neuron *j* of type *X* and *Y* (excitatory or inhibitory) evolved such that:

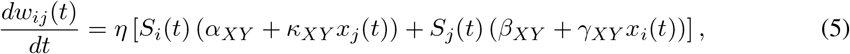

with *η* = 0.01 a fixed learning rate, 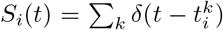 the spike train of neuron *i, δ* the Dirac delta function to denote the presence of a pre (post)-synaptic spike at time *t*. The synaptic traces *x*_*i*_ and *x*_*j*_ are low-pass filters of the activity of presynaptic neuron *i* and postsynaptic neuron *j*, with time constants *τ*_pre_ and *τ*_post_, such that:

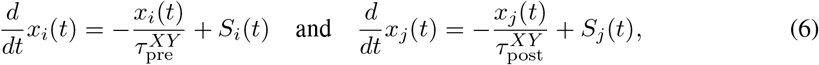

Overall, this search space comprised 6 tunable plasticity parameters per synapse type 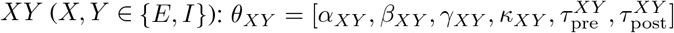. The four synapse-type vectors together give 24 parameters per rule. In this work we do not search this space de novo: we use the library of 2201 rules released with prior work [13], each obtained there via simulation-based inference and retained for producing stable network dynamics.

Note that all weights in the network were capped at all times, in the [0, *w*_max_] range, with *w*_max_ = 20, though rule quadruplets in [21] were considered unstable if more than 10% of the weights at any synapse type reached these extreme values.

### A.2 Spiking reservoir computing setup in Pong framework

The Pong framework was adapted from prior work [24, 13]. The game environment was rendered on a 64 × 64 grid, each corresponding to one excitatory neuron in the reservoir. At each timestep, the ball position was translated into a 2D grid coordinate (*r, c*) and communicated to the spiking network server. The network was simulated for 200 ms per timestep using the Auryn simulator [51], returning spike counts for all *N*_*E*_ = 4096 excitatory and *N*_*I*_ = 1024 inhibitory neurons.

The spatial stimulus was encoded as follows. At each timestep, the ball position (*c, r*) on the 64 × 64 grid was used to activate a 9 × 9 patch of excitatory neurons centered on the corresponding grid location. The 81 neurons within the patch received Poisson input at rate *r*_stim_ = 100 Hz for the full 200 ms simulation window, while all other excitatory neurons received no direct stimulus drive. The patch boundaries were clipped to the grid edges to handle ball positions near the boundary. This stimulus group is separate from the background Poisson population (*N*_inp_ = 10,000, *r*_bg_ = 10 Hz), which provided ongoing drive to both excitatory and inhibitory neurons via receptive-field connections throughout the simulation.

At each such timestep, the spike rates of all excitatory neurons were passed to the linear readout together with the ball’s current position and its prior position (i.e. **r**_exc_(*t*) decoded against (*x*_0_, *y*_0_, *x*_*−*1_, *y*_*−*1_)). Firing rates were normalized to [0, 1] before being passed to the readout. Ball positions were normalized to [−1, 1].

The linear readout comprised a single fully connected layer mapping *N*_*E*_ = 4096 excitatory neuron firing rates to four output units predicting the current 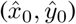 and a past 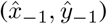 ball coordinates. The readout was trained with the Adam optimiser (learning rate 10^*−*3^) using mean squared error loss, summed over current and past position predictions:

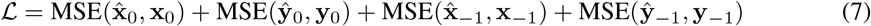

At the end of each game, the readout was updated on all samples accumulated during that game, after which the buffer was cleared. During testing, readout weights were frozen and loss was computed without gradient updates.

The paddle position was predicted by fitting a line through the readout’s predicted current and past ball positions using least squares, and extrapolating to the left wall (*x* = −1). The predicted *y*-coordinate was clipped to [−1, 1] and rescaled to screen coordinates.

Training proceeded in three phases (Fig. 1C). First, a burn-in period of approximately one minute with plasticity on allowed the network to reach a stable activity regime. This was followed by 1000 training games with plasticity on and readout learning on (online). Finally, 250 testing games were run with plasticity on and readout learning off. Ball direction was randomly initialized at the start of each game, and ball velocity could change upon wall contact, making direction prediction non-trivial (Fig. 1B). Plasticity was kept on during testing to reflect a continuous online learning scenario; Supplementary Figure S4 confirms that freezing the reservoir after training shows comparable performance, validating that the learned connectivity is the primary driver of test performance.

### A.3 Spiking reservoir computing setup in Delayed-match (DM) framework

The delayed-match (DM) task was implemented on the same recurrent spiking reservoir and plasticity parameterization as the Pong framework. The network was simulated in 200 ms timesteps using the Auryn simulator [51], returning spike counts for all *N*_*E*_ = 4096 excitatory and *N*_*I*_ = 1024 inhibitory neurons.

The stimulus set comprised a bank of 13 fixed assemblies *S*_0_, …, *S*_12_, each a disjoint group of 300 excitatory neurons selected at initialization and held constant across trials. On each trial, two patterns were drawn from the bank and presented sequentially: a sample pattern for *t*_stim_ = 1 s, a delay of *t*_delay_ = 5 s during which no assembly was driven, and a test pattern for *t*_stim_ = 1 s. During each stimulus window the 300 neurons of the active assembly received Poisson input at rate *r*_stim_ = 100 Hz. This stimulus group is separate from the background Poisson population (*N*_inp_ = 10,000, *r*_bg_ = 10 Hz), which provided ongoing drive to both excitatory and inhibitory neurons via receptive-field connections throughout the trial. A trial was labelled “match” if the sample and test patterns were identical and “no-match” otherwise, with match and no-match trials balanced and no-match test patterns drawn uniformly from the remaining assemblies.

At each timestep, the spike rates of all excitatory neurons were normalized to [0, 1] and passed to a linear readout. The readout comprised a single fully connected layer mapping the *N*_*E*_ = 4096 excitatory firing rates to a match/no-match decision, trained with the Adam optimizer (learning rate 10^*−*3^) under a binary cross-entropy loss. At the end of each training trial, the readout was updated on the samples accumulated during that trial, after which the buffer was cleared; during testing the readout weights were frozen and no gradient updates were applied.

Training proceeded in the same three phases. First, a burn-in period of one minute with plasticity on allowed the network to reach a stable activity regime. This was followed by 1000 training trials with plasticity on and readout learning on (online), and finally 250 testing trials with plasticity on and readout learning off. Trial-level classification accuracy on the held-out testing trials was used as the DM performance measure (chance = 0.5).

### A.4 Analysis of activity and weights

#### A.4.1 Pairwise cosine similarity of the readout units

The linear readout consisted of four units, each predicting one coordinate of the ball’s current or previous position (*y*_0_, *x*_0_, *y*_*−*1_, *x*_*−*1_). The readout weight vectors were extracted from the trained readout after the testing phase. We computed pairwise cosine similarities between all six pairs of weight vectors:

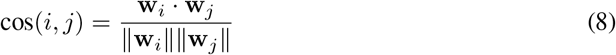

Cosine similarities were computed per seed and averaged across seeds. The mean pairwise cosine similarity across all six pairs was used as a summary alignment statistic. Spearman correlations between individual pair cosine similarities and average testing score were computed across all *n* = 2201 networks.

#### A.4.2 Mean firing rate of ball trajectory and background neurons

Activity of excitatory neurons was analyzed from 200 ms binned spike counts recorded during the testing phase. For each timestep, neurons were assigned to one of groups: the 9×9 neurons centered on the current ball position (*t*_0_), and the three past ball positions (*t*_*−* 1_, *t* _*−*2_, *t*_*−* 3_), with all remaining neurons constituting the background (bg.). Spike counts were converted to firing rates and the median rate across timesteps was computed. Per-frame normalization was applied before computing zone averages, allowing rules with different overall activity levels to be compared directly.

#### A.4.3 Weights statistics and drift

Synaptic weight matrices were extracted at two time points: the end of training and the end of testing phase. For each of the four connection types (EE, EI, IE, II), mean weight 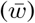 and standard deviation (*σ*_*w*_) were computed from the non-zero weight values at the end of training. Weight drift between training and testing phases was quantified as the mean absolute change ⟨| Δ*w*|⟩ across all synapses of a given connection type.

### A.5 Regression models

All regression models used Ridge regression with the regularization strength *α* selected by 5-fold cross-validation over a log-spaced grid (*α* ∈ [10^*−*3^, 10^5^], 100 values). Performance was evaluated by 5-fold cross-validated *R*^2^ and RMSE on the full dataset of *n* = 2201 networks. All feature matrices were standardized to zero mean and unit variance prior to fitting.

#### A.5.1 Parametrization of the rules as predictors

In Tier 1, we used the 24 plasticity rule parameters directly as predictors: *α*_*XY*_, *β*_*XY*_, *γ*_*XY*_, *κ*_*XY*_, 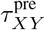, and 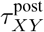 for each synapse type (*X, Y*) ∈ {*EE, EI, IE, II*} (*k* = 24). This model is computation-free. It required no simulation beyond what was already available from the rule parameterization itself.

#### A.5.2 Functionality of the rules in memory tasks and the initial slope of the loss

For Tier 2a, we derived performance on the novelty detection task as predictors. The task was implemented in the prior work [13]. For each plasticity rule, the absolute change in response magnitude | Δ*r*_mem_| was computed across 10 break durations (*t*_break_ ∈ {1, 10, 20, 60, 120, 300, 600, 1200, 3600, 14400} s), yielding 10 features (*k* = 10) (see also Supplementary Figure S5 for correlation of Pong performance with other memory tasks).

For Tier 2b, we used the initial slope of the training loss (ℒ) as a single predictor (*k* = 1). The slope was estimated as the coefficient of a linear fit of log_10_(ℒ) against game number over the first 200 training games.

#### A.5.3 After-training features as predictors

##### Tier 3a — Readout alignment (*k* = 12)

We computed pairwise cosine similarities between all six pairs of readout weight vectors, summarized as four statistics: the mean cosine similarity across cross pairs, same-coordinate pairs, same-time pairs, and all six pairs combined. Additionally, we included six L2-norm features: the norm of each of the four readout weight vectors individually, their mean, standard deviation, and the ratio of the mean norm of current-position units to past-position units (∥**w**_now_∥ */* ∥**w**_past_∥), together with the variance explained by the first principal component of the readout weight matrix, for a total of 12 features.

##### Tier 3b — Activity contrast (*k* = 9)

We used the mean firing rate of excitatory neurons in each group across all testing timesteps: rates at the current and three preceding ball positions (*t*_0_, *t*_*−*1_, *t*_*−*2_, *t*_*−*3_) and the background rate (bg.), yielding 5 rate features. We additionally included four derived contrast features: the ratio of mean stimulus activity to background, 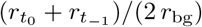, quantifying overall stimulus detectability; the relative decay from current to past, 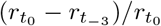, quantifying the sharpness of the temporal memory trace; the mean firing rate across all four stimulus zones; and the ratio of the current-position rate to background, 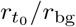, for a total of 9 features.

##### Tier 3c — Reservoir weights (*k* = 20)

For each connection type, we extracted three statistics of the weight distribution after training (mean 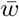, standard deviation *σ*_*w*_, and coefficient of variation 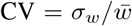 and two drift statistics between the end of training and end of testing: the absolute mean weight change ⟨|Δ*w*|⟩ and the Pearson correlation of weight vectors before and after testing, yielding 20 features.

### A.6 Benchmark models

All benchmark models were embedded in the same Pong framework: identical 64 × 64 spatial input encoding, same three-phase protocol (burn-in, 1000 training games, 250 testing games). All models were evaluated across multiple seeds (from two to six).

#### A.6.1 Regression

The regression model used no reservoir. The normalized ball position (*x, y*) ∈ [− 1, 1]^2^ was passed directly to the linear readout as a 2-dimensional input vector, with Gaussian noise N(0, *σ*^2^) added to the input to prevent trivial memorization. Six noise levels were evaluated (*σ* ∈ {0, 0.1, 0.325, 0.55, 0.775, 1.0}), in total 6 configurations.

#### A.6.2 Static network

The static network used the same LIF reservoir as the plastic models (see Network model), with plasticity disabled throughout (*η* = 0). To vary the representational capacity of the reservoir, the initial synaptic weights were drawn from a log-normal distribution with fixed mean (*w*_*EE*_ = *w*_*EI*_ = 0.1, *w*_*IE*_ = *w*_*II*_ = 1.0) and varying standard deviation *σ*, controlling the degree of weight heterogeneity. At *σ* = 0 all weights were initialized homogeneously to their mean values. Six heterogeneity levels were evaluated (*σ* ∈ {0, 0.01, 0.0325, 0.055, 0.0775, 0.1}), in total 6 configurations. The readout received the spike counts of the *N*_*E*_ = 4096 excitatory neurons, normalized to [0, 1].

#### A.6.3 Static metabolic network

The static metabolic network replaced the standard LIF neurons with metabolic integrate-and-fire (MIF) neurons [28], simulated using the Auryn simulator [51]. MIF neurons incorporate a metabolic adaptation variable *A*(*t*) with slow and fast timescales (*τ*_*A*_ = 300 ms, 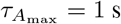, *A*_max_ = 25 mV) and a firing threshold that depends on local metabolic state via parameters *K*_*µ*_ and *K*_*σ*_. Synaptic weights were static and optionally initialized from a log-normal distribution to introduce heterogeneity, with mean weights *w*_*EE*_ = *w*_*EI*_ = 0.2 and *w*_*IE*_ = *w*_*II*_ = 3.0. The swept hyperparameters were *K*_*µ*_ ∈ {0.2, 0.25, 0.3, 0.35, 0.4, 0.45, 0.5} and coefficient of variation CV ∈ {0.0, 0.05, 0.10}, giving 21 configurations. The spatial stimulus and readout interface were identical to the static LIF network. Unlike the static LIF network, no background Poisson input was provided; the network received only the spatial stimulus drive.

#### A.6.4 ESN

Echo state networks [29–31] used a rate-coded reservoir with sparse random connectivity (10%). For Dale’s law configurations, the reservoir comprised *N*_*E*_ = 4096 excitatory and *N*_*I*_ = 1024 inhibitory units with sign-constrained weights; for non-Dale configurations, a single recurrent population of *N* = 5120 units with unconstrained weights was used. At each timestep, the ball position (*x, y*) normalised to [−1, 1]^2^ was passed as a 2-dimensional input to the reservoir, scaled by the input scaling parameter *σ*_in_. The reservoir state was integrated for *T* discrete steps using leaky integration with rate *γ*_leak_, and the resulting state vector was passed to the linear readout. The swept hyperparameters were spectral radius *ρ* ∈ [0.53, 1.49], leak rate *γ*_leak_ ∈ [0.09, 1.0], input scaling *σ*_in_ ∈ [0.06, 2.92], and integration steps *T* ∈ {10, 15, 20, 25, 30}, yielding 21 Dale and 29 non-Dale configurations (50 total).

#### A.6.5 FORCE

FORCE learning [32] used the same rate-coded reservoir as the ESN (*N*_*E*_ = 4096, *N*_*I*_ = 1024, 10% connectivity) with an additional feedback connection from the readout output back into the reservoir, scaled by *g*_fb_. The readout weights were trained online using recursive least squares (RLS) with regularization *α*_RLS_, updated at every timestep during training. The ball position was passed as a 2-dimensional input scaled by *σ*_in_, and the reservoir was integrated for *T* steps with spectral radius *ρ* and leak rate *γ*_leak_. The swept hyperparameters were *ρ* ∈ [0.80, 1.46], *γ*_leak_ ∈ [0.10, 0.72], *σ*_in_ ∈ [0.11, 1.97], *g*_fb_ ∈ [0.74, 2.97], *α*_RLS_ ∈ [0.91, 8.95], and *T* ∈ {10, 15, 20, 25}, yielding 20 configurations.

#### A.6.6 LSTM Pred

The LSTM predictive model [33, 34] used a multi-layer LSTM network as the reservoir replacement. The normalized ball position (*x, y*) ∈ [− 1, 1]^2^ was passed as a 2-dimensional input at each timestep to a stacked LSTM with hidden size *h* and *L* layers, followed by a linear projection to *N*_*E*_ = 4096 output dimensions. The output was passed to the same linear readout as all other models. The LSTM was trained with a self-supervised predictive objective: at each timestep, the network predicted the next ball position, and the prediction error was used to update the LSTM weights via Adam optimizer. The linear readout was trained separately on the LSTM’s output representations, identical to all other models. The hyperparameters hidden size *h* ∈ {128, 256, 512}, number of layers *L* ∈ {1, 2, 3}, and learning rate *η* ∈ [10^*−*4^, 3 × 10^*−*3^] were explored via random search over 20 configurations.

#### A.6.7 e-prop

The e-prop model [35] was implemented as a two-layer spiking network: a recurrent LIF core of *n*_rec_ neurons (sparse, 10% connectivity, Dale-constrained with 80% excitatory) projecting to *N*_*E*_ = 4096 LIF output neurons via a sparse output weight matrix *W*_out_. The ball position (*x, y*) normalised to [−1, 1]^2^ was projected into the recurrent core via a dense input layer 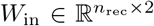. At each outer timestep, the network was unrolled for *T* inner LIF steps; the resulting output spike counts were passed to the linear readout. The recurrent and output weights were updated online at each outer step using random e-prop: eligibility traces 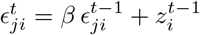 were maintained for all plastic synapses, and weight updates were computed as Δ*W*∝ *L*_*j*_ ·*e*_*ji*_, where *L*_*j*_ = *B*_*j*_ ·*δ*_out_ is the learning signal propagated via fixed random feedback weights *B* from the output error *δ*_out_, and *e*_*ji*_ = *ϵ*_*ji*_ ·*ψ*_*j*_ with *ψ*_*j*_ the pseudoderivative damped by *γ*. Memory was *O*(*W*) independent of game length. An internal supervised head 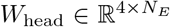 provided the output error signal, trained on the same 4-dimensional ball-position target as the external readout. The swept hyperparameters were *n*_rec_ ∈ {256, 512, 768}, inner steps *T* ∈ {10, 15, 20, 25}, membrane decay *β* ∈ [0.73, 0.93], pseudoderivative damping *γ* = 0.3 (fixed), and learning rate *η* ∈ [10^*−*4^, 2.5 × 10^*−*3^], yielding 20 configurations.

#### A.6.8 SNN — surrogate gradient

Two surrogate-gradient SNN variants were evaluated, both sharing the same two-layer LIF architecture implemented using snnTorch [58]: a recurrent LIF core of *n*_rec_ neurons (sparse, 10%, Dale-constrained, 80% excitatory) projecting to *N*_*E*_ = 4096 LIF output neurons via sparse *W*_out_. The ball position was projected into the core via a dense input layer. At each outer timestep the network was unrolled for *T* inner LIF steps, with LIF membrane decay *β* initialised to a swept value and set as a learnable parameter. Output spike counts were passed to the linear readout. Hidden state (membrane potential and spikes) was maintained persistently across timesteps within a game and reset at game boundaries.

In the **end-to-end** (E2E) variant, an internal supervised head 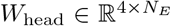 received the same 4-dimensional ball-position target as the external readout. Gradients were propagated through the surrogate derivative of the spike function back through the full two-layer network using truncated BPTT, flushed every 100 outer steps to bound memory usage. All weights (*W*_in_, *W*_rec_, *W*_out_, *W*_head_, *β*) were updated via Adam at every flush.

In the **predictive** (Pred.) variant, no task-level error signal reached the recurrent network. Instead, a small predictive head (Linear(*N*_*E*_, 64)–ReLU–Linear(64, 2)) was trained to predict the next normalised ball position from the current output spike counts, updated at every outer step via per-step BPTT with hidden state detached after each update. The recurrent network weights were updated by this self-supervised predictive loss alone; the external readout was trained identically to all other models.

For both variants, swept hyperparameters were *n*_rec_ ∈ {256, 512, 768}, inner steps *T* ∈ {10, 15, 20, 25}, membrane decay *β* ∈ [0.71, 0.94], and learning rate *η* ∈ [10^*−*4^, 4 ×10^*−*3^], yielding 20 configurations.

### A.7 Multi-fidelity SBI for sampling Pong expert rules

#### A.7.1 Mean-field approximation and surrogate score function

Following the mean-field model of prior work [21], the steady-state firing rates satisfied four coupled equations, one per each connection type:

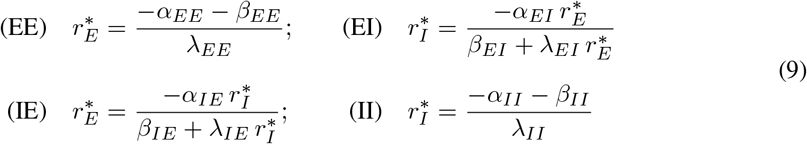

with 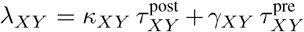 for (*X, Y*) ∈ {*E, I*}. We computed *λ*_*XY*_ for all four connection types and obtained steady-state rates from the (EE) and (EI) equations, giving 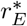 directly from (EE) and 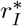 from (EI) given 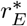. A rule was classified as mean-field stable if 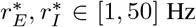, consistent with prior work [22].

Candidate rules were sampled uniformly from the prior (*τ*_*XY*_ ∈ [0.01, 0.1] s; *α*_*XY*_, *β*_*XY*_, *γ*_*XY*_, *κ*_*XY*_ ∈ [−2, 2])in batches of 2 × 10^6^, retaining those passing the rate-range filter until *N*_LF_ = 100,000 stable low-fidelity samples were collected.

To assign Pong scores to these low-fidelity samples, we trained a surrogate score function on the *N*_HF_ = 2201 high-fidelity rules. We compared a *k*-nearest-neighbour regressor (with *k* selected by 5-fold cross-validation over *k* ∈ {3, 5, 10, 15, 20, 30, 50}) against a histogram-based gradient boosting regressor (500 iterations, learning rate 0.05, max depth 6). Cross-validated performance on the high-fidelity set favoured the gradient boosting regressor (CV *R*^2^ = 0.346 vs. 0.201 for *k*-NN, compared to a predict-mean baseline of *R*^2^ =− 0.002), and it was therefore selected as the surrogate. Each low-fidelity rule was assigned a surrogate-predicted pseudo-label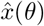.

#### A.7.2 MF-SBI pipeline

We proceeded in two stages (Fig. 6A), following [22]. First, a Neural Posterior Estimator (NPE) was pretrained on the *N*_LF_ = 100,000 low-fidelity 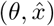 pairs, with a batch size of 512, learning rate 5×10^*−*4^. In the second stage, the pretrained NPE was used to initialize a second NPE, which was then fine-tuned on the *N*_HF_ = 2201 high-fidelity samples using the same architecture and training procedure.

We sampled 5,000 rules from the resulting posterior *π*_1_ under two conditioning observations: the mean score of the top 1% of screened rules (*x*_*o*_ = 3.72) and the maximum high-fidelity testing score (*x*_*o*_ = 5.14). Posterior samples are not guaranteed to produce stable activity and weight dynamics. We therefore applied a post-hoc mean-field rate filter before submitting rules to full spiking network evaluation: a sampled rule was retained only if 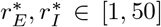 Hz under Eqs. 9. Of the 5,000 samples drawn per conditioning value, 934 (*x*_*o*_ = 3.72), and 900 (*x*_*o*_ = 5.14) passed this filter and were forwarded for evaluation in the Pong framework.

#### A.7.3 Filter for unstable samples - post-simulation

Following full evaluation in the Pong framework, we applied a second post-hoc stability filter to remove rules that produced unstable network dynamics during simulation. Specifically, a rule was excluded if (i) the mean firing rate of excitatory or inhibitory neurons fell outside the range [1, 50] Hz, or (ii) the fraction of weights reaching the maximum synaptic cap (*w*_max_ = 20 nS) exceeded 20% for any synapse type at the end of training. After applying this filter and simulating across three seeds, 615 (*x*_*o*_ = 3.72) and 489 (*x*_*o*_ = 5.14) rules remained and were used for all subsequent analyses.

#### A.7.4 Held-out seed re-scoring

To test whether posterior enrichment depends on the seeds used to condition the SBI, we re-evaluated rules on three held-out seeds disjoint from screening and inference. Re-scoring the full 2201-rule prior is as costly as the original screen, so we scored a subsample of prior rules stratified by original-seed score and reweighted each stratum by its Horvitz–Thompson weight *w*_*h*_ = *N*_*h*_*/n*_*h*_. The reweighted subsample reproduces the full-prior distribution on the original seeds (Fig. S2A) and per-rule scores are preserved across seed sets (Fig. S2B).

**Table S1:** Regression models predicting average testing score from plasticity rules parametrization.

| Regression model | *k* | CV *r*<sup>2</sup> | CV RMSE | Spearman *ρ* |
|---|---|---|---|---|
| Ridge regression (linear) | 24 | 0.352 | 0.549 | 0.66 |
| Multi-layer perceptron (MLP) | 24 | 0.328 | 0.559 | 0.64 |
| **TabPFN** | 24 | **0.557** | **0.454** | **0.79** |

**Table S2:**
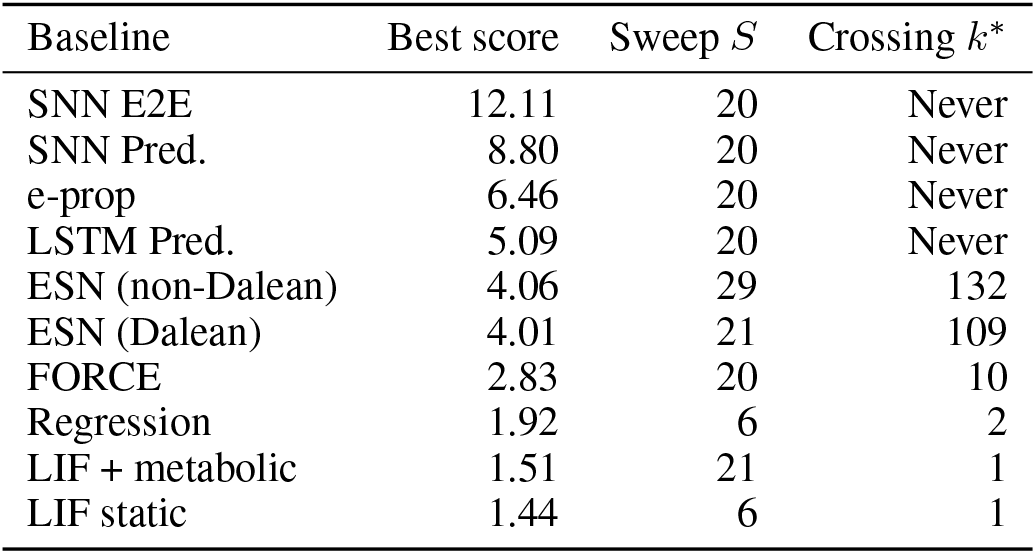
Budget-normalised benchmark comparison. For each baseline we report its best score and the plastic search budget *k*^*∗*^ at which the expected best-of-*k* plastic rule matches it. *k*^*∗*^ is the expected maximum over random subsamples of the screened plastic pool (*n* = 2201). “Never” indicates the baseline is not overtaken within the screened pool.

## A.8 Supplementary figures

**Figure S1:**
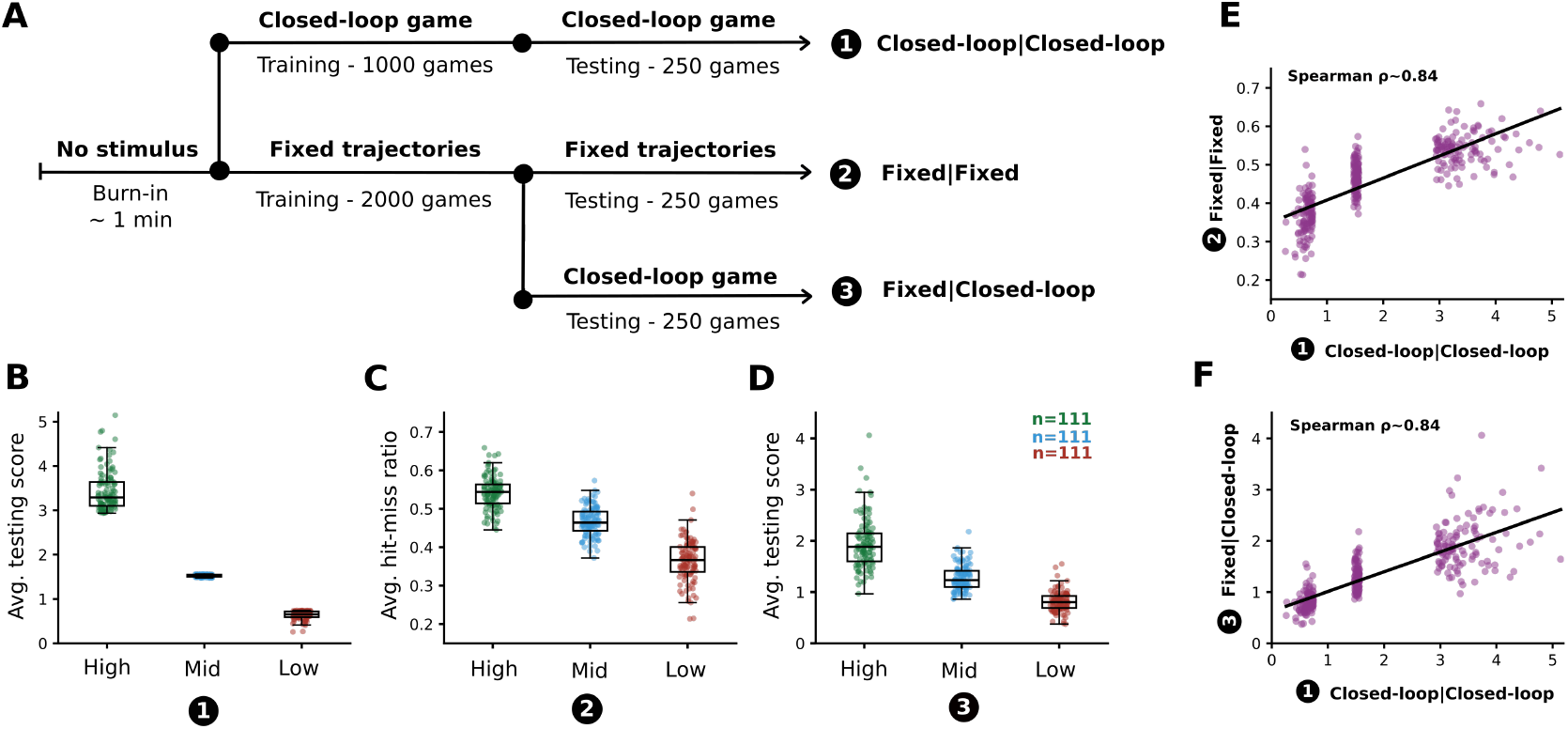
Fixed-trajectory exposure control disentangles rule quality from training exposure. **A**. Schematic of the three testing protocols compared, following a shared no-stimulus burn-in (~1 min). (1) Closed-loop | Closed-loop: the original protocol (1000 training games, 250 testing games), subject to the Matthew effect. (2) Fixed | Fixed: reservoir and readout trained (2000 games) and tested (250 games) on fixed-length trajectories that terminate as soon as the ball reaches the paddle or wall, equalizing exposure across rules; scored by hit-miss ratio rather than average score. (3) Fixed|Closed-loop: reservoir and readout trained on fixed trajectories (2000 games), then tested in the original closed-loop setting (250 games). **B**. Average testing score under protocol (1) for high-, mid-, and low-scoring rule groups (*n* = 111 each, the top/bottom/middle 5% of screened rules). **C**. Average hit-miss ratio under protocol (2) for the same three groups. **D**. Average testing score under protocol (3) for the same three groups (*n* = 111 per group). **E**. Per-rule correlation between scores under protocols (2) and (1). **F**. Per-rule correlation between scores under protocols (3) and (1).

**Figure S2:**
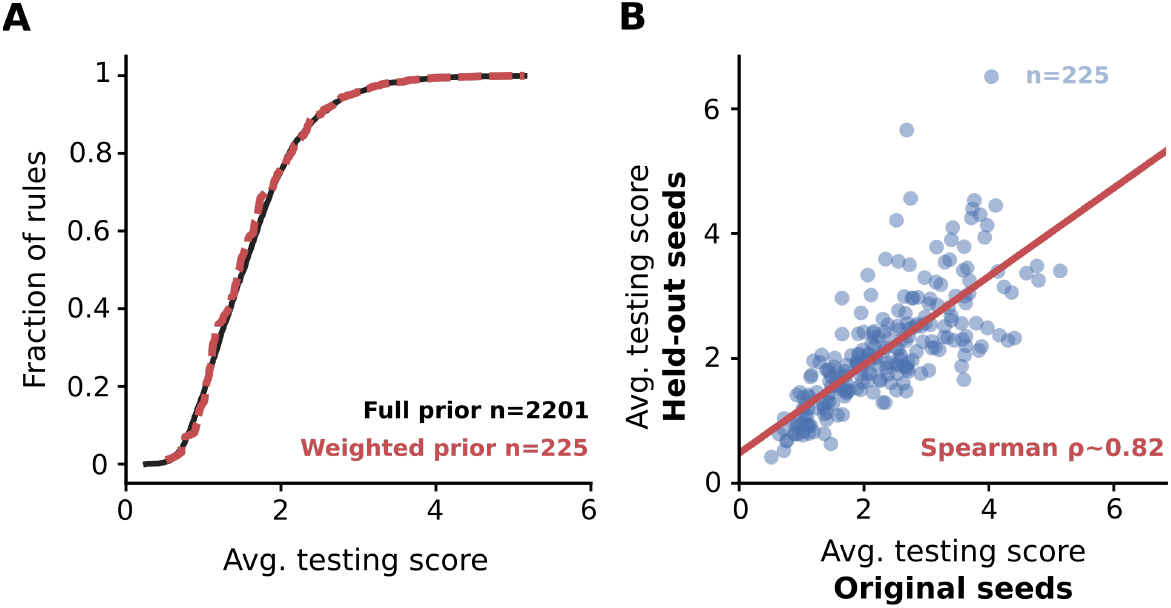
All rules re-scored on three held-out seeds not used during screening or SBI. **A**. The Horvitz–Thompson-reweighted subsample (red) reproducing the full prior (black) on the original seeds. **B**. Correlation of per-rule scores are preserved between original and held-out seeds (Spearman *ρ* = 0.82), consistent with the across-seed variability in Fig. S7.

**Figure S3:**
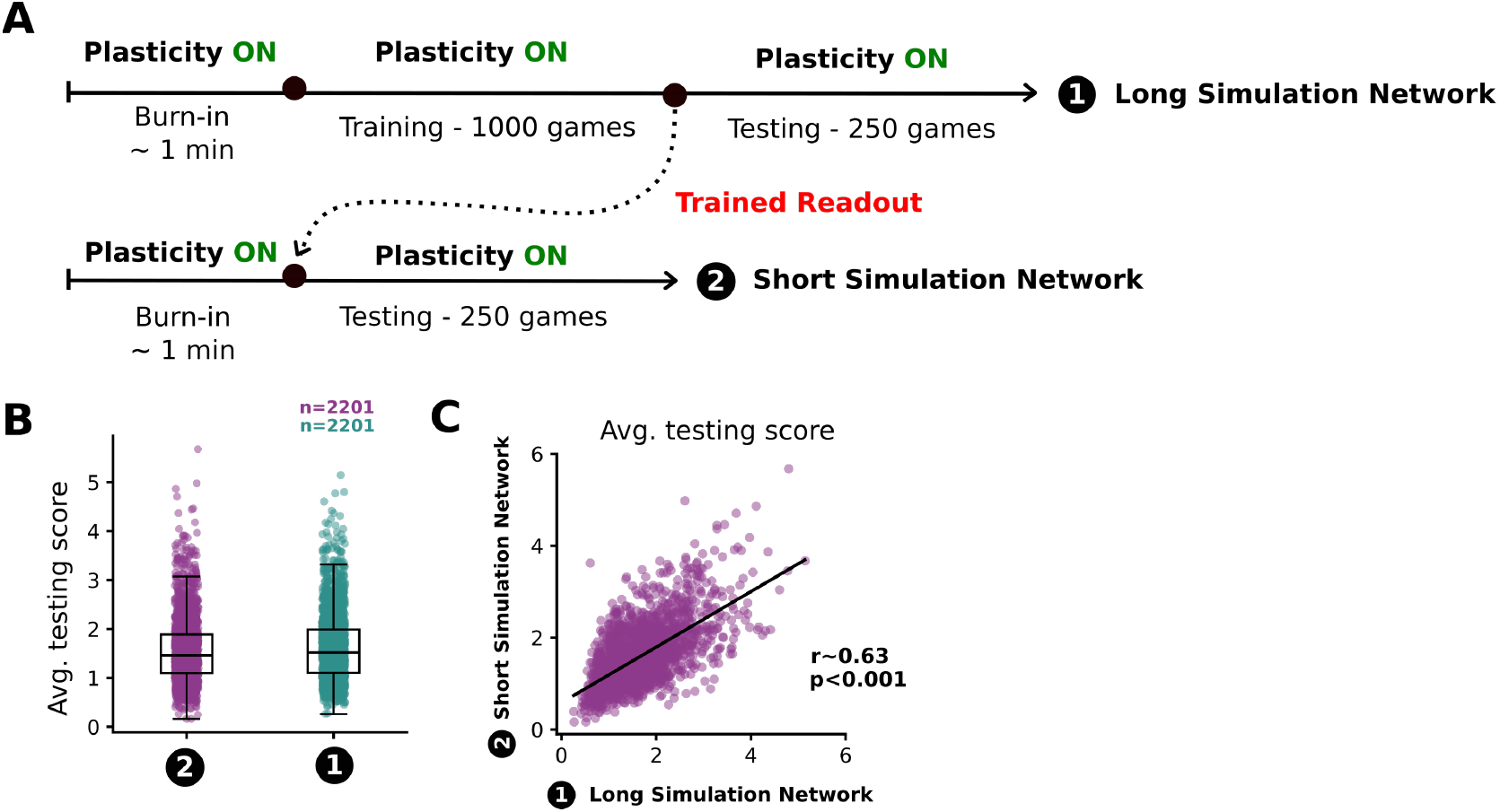
Transfer of a trained readout to a new network. **A**. Schematic of the two network conditions. The long simulation network undergoes burn-in, training (1000 games), and testing (250 games), all with plasticity ON. A trained readout is then transferred to the short simulation network, which undergoes only burn-in and testing (250 games) with plasticity ON. **B**. Average testing scores for the short simulation network (purple) and long simulation network (teal); *n* = 2201 per condition. **C**. Correlation of average testing scores between the two networks.

**Figure S4:**
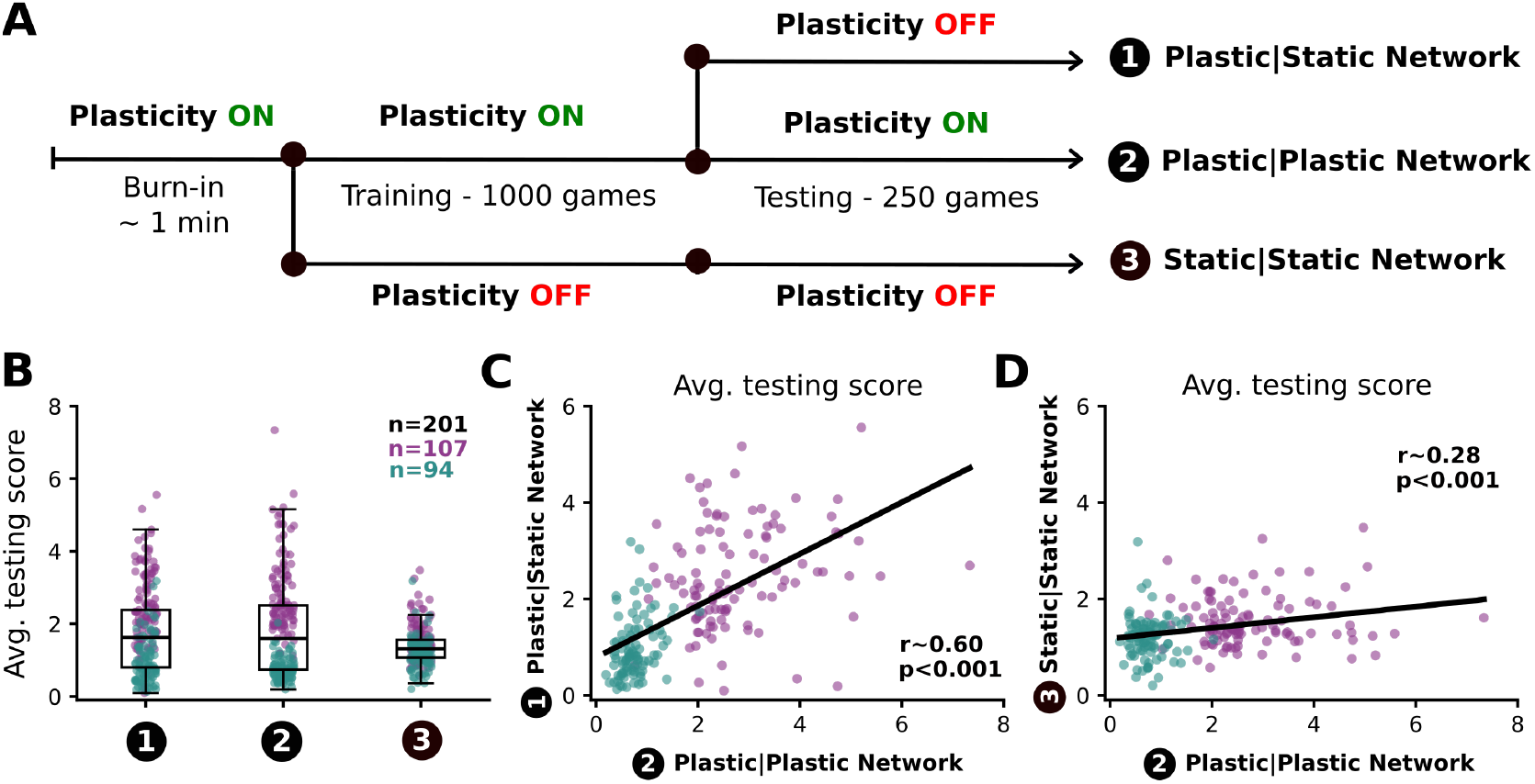
Effect of static vs plastic reservoirs during and after training. **A**. Schematic of the three network conditions. All networks undergo burn-in and training (1000 games) with plasticity ON, after which they diverge: the Plastic|Static network switches plasticity OFF during testing, the Plastic|Plastic network maintains plasticity ON throughout testing, and the Static|Static network has plasticity OFF during both training and testing. All conditions are tested over 250 games. **B**. Distribution of average testing scores across the three conditions for the same *n* = 201 networks evaluated under Plastic|Static, Plastic|Plastic, and Static|Static settings, with 107 high-scoring (purple) and 94 low-scoring (teal) networks. This subset was selected as the union of the top and bottom 5% of screened rules by average testing score. **C**. Correlation of average testing scores of the Plastic|Static versus Plastic|Plastic networks. **D**. Correlation of average testing scores of the Static|Static versus Plastic|Plastic networks.

**Figure S5:**
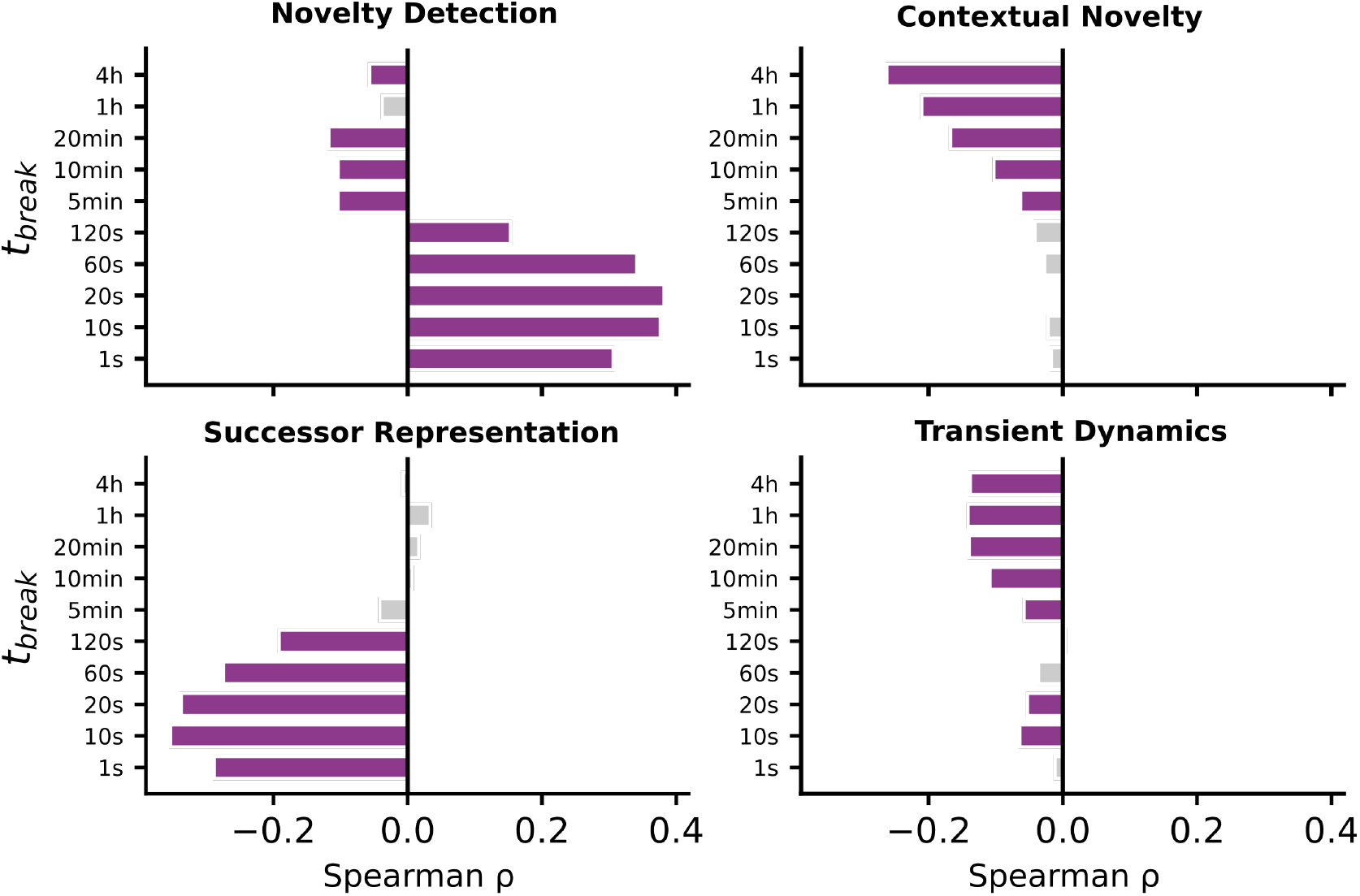
Correlation between memory task performance and average testing score across break durations in a task. Spearman correlation (*ρ*) between average testing score and performance on each of the four memory tasks from prior work [13] computed separately for each break duration (*t*_break_) in the task. Purple bars indicate statistically significant correlations; grey bars indicate nonsignificant results.

**Figure S6:**
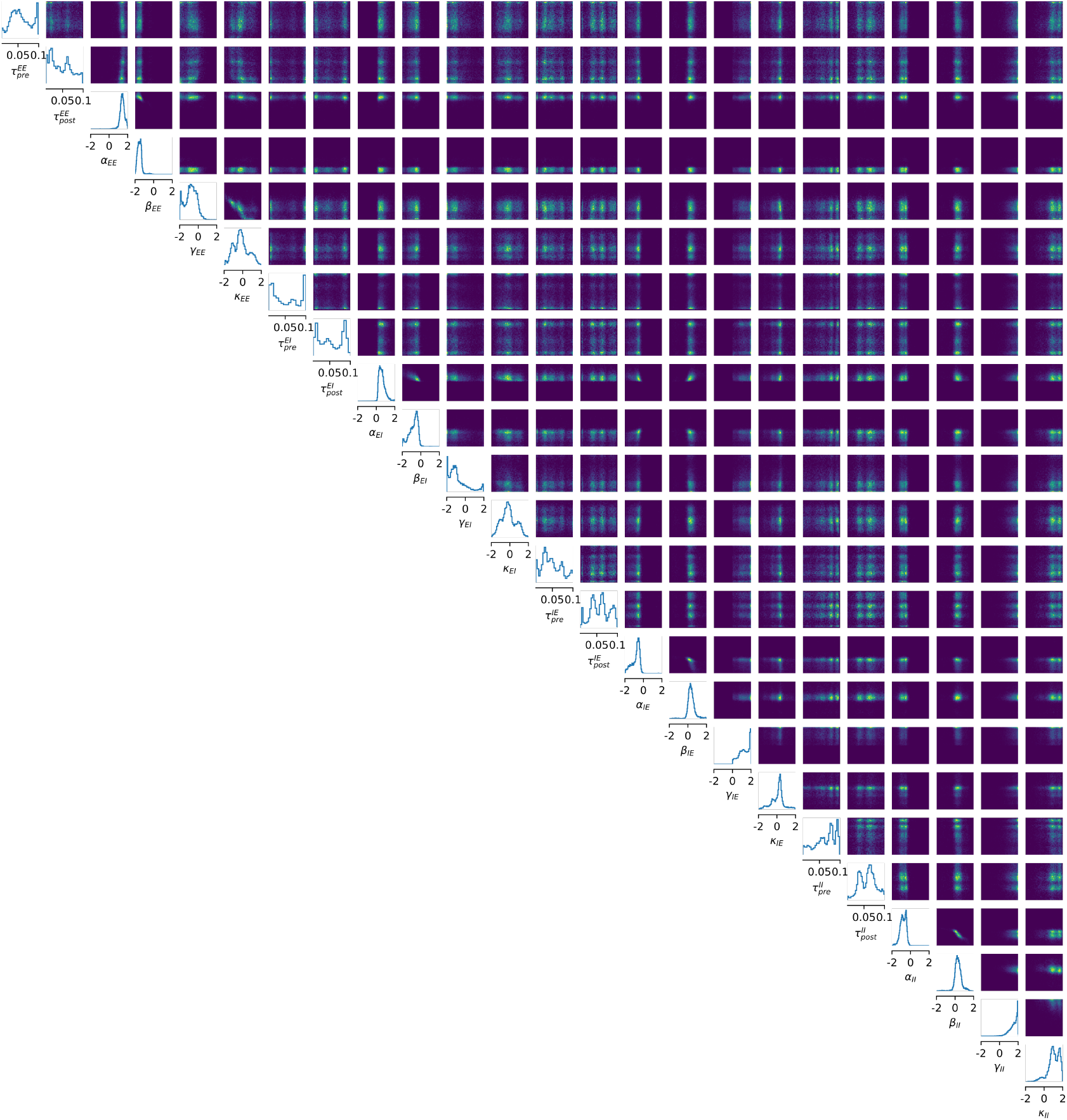
Visualization of posterior for *x*_*o*_ = 5.14. Pairplot of 10,000 samples from the posterior *π*_1_. The diagonal panels show the distribution of values for each plasticity parameter, while the off-diagonal panels show the marginal distributions between each pair of plasticity parameters.

**Figure S7:**
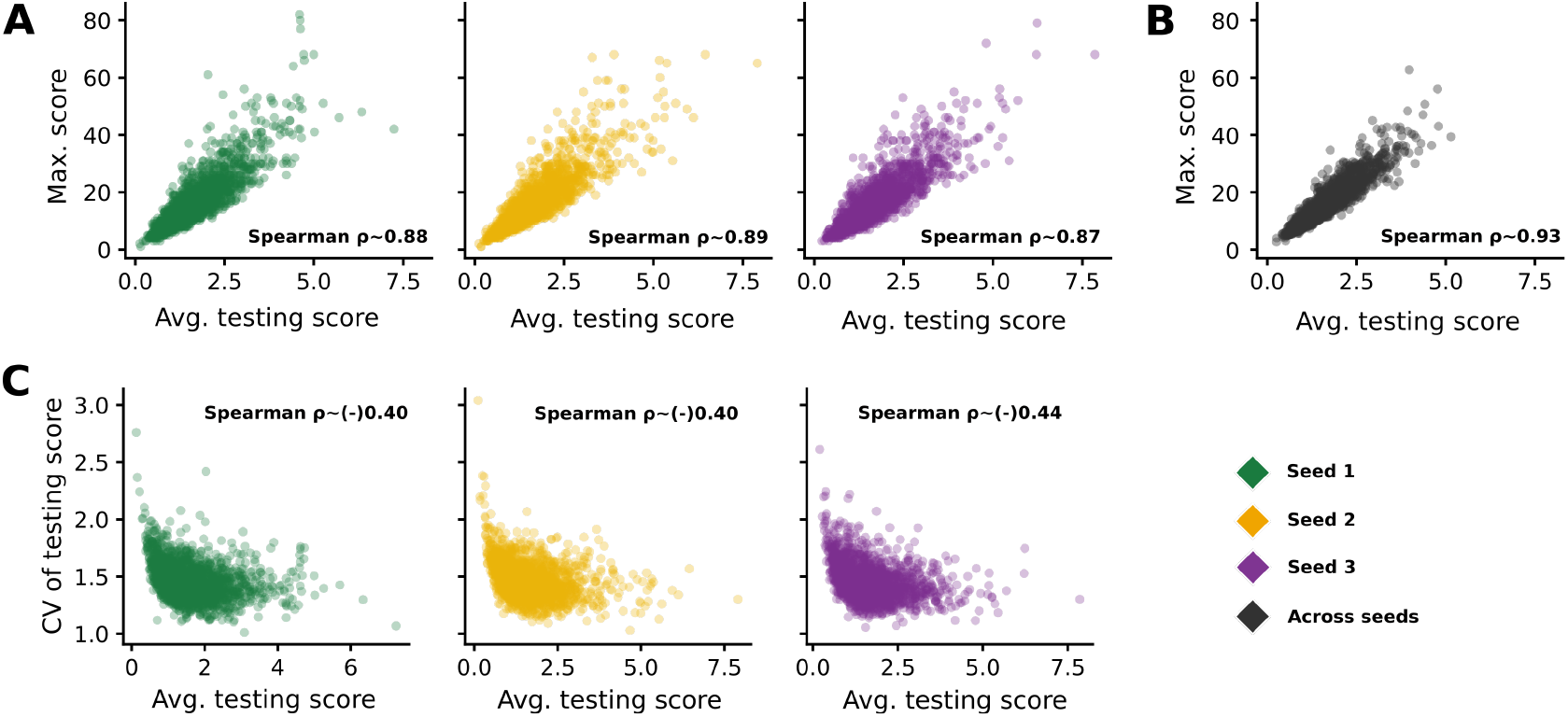
Rule consistency: maximum, average, and variability of testing scores. Each point is a single plasticity rule (*n* = 2201). **A**. Maximum vs. average testing score per seed (Spearman *ρ* ~ 0.88, 0.89, 0.87). **B**. The same relationship with scores averaged across the three seeds (Spearman *ρ* ~ 0.93). **C**. Coefficient of variation (CV) of the testing score vs. average testing score per seed (Spearman *ρ* ~ −0.40, −0.40, −0.44).

